# An anti-virulence drug targeting the evolvability protein Mfd protects against infections with antimicrobial resistant ESKAPE pathogens

**DOI:** 10.1101/2024.01.22.576688

**Authors:** SL. Tran, L. Lebreuilly, D. Cormontagne, S. Samson, TB. Tô, R. Dervyn, A. Grießhammer, J. de la Cuesta-Zuluaga, L. Maier, T. Naas, S. Mura, J. Nicolas, D. Rognan, G. André, N. Ramarao

## Abstract

The increased incidence of antibiotic resistance and declining discovery of new antibiotics have created a global health crisis, especially for the treatment of infections caused by Gram-negative bacteria. Here, we identify and characterize a molecule, NM102, that displays antimicrobial activity exclusively in the context of infection. NM102 inhibits the activity of the non-essential Mutation Frequency Decline (Mfd) protein by competing with ATP binding to its active site. Inhibition of Mfd by NM102 sensitizes pathogenic bacteria to the host immune response and blocks infections with clinically- relevant *Klebsiella pneumoniae* and *Pseudomonas aeruginosa*, without inducing host toxicity. Finally, NM102 inhibits the function of Mfd as a mutation and evolvability factor, thus reducing the bacterial capacity to develop antimicrobial resistance. These data provide a potential roadmap to expand the arsenal of drugs to combat antimicrobial resistance.

**Highlight:**

- NM102 is a “first in class” molecule specifically targeting the active site of the bacterial Mfd protein
- NM102 has a new mode of action: it inhibits Mfd function during immune stress response
- NM102 also inhibits Mfd evolvability function and thereby decreases bacterial resistance to known antibiotics
- NM102 effectively treats Gram-negative infections in animal models
- NM102 is efficient against clinically relevant resistant bacteria and provides an increased efficacy in combination with the β-lactam meropenem

## Introduction

The fight against pathogenic bacteria is turning again into one of the greatest challenges for our societies, especially with the spread of multi- and extensively-drug resistant bacteria, which seriously imperil the use of traditional antibiotics (Cook and Wright, 2022). Estimates suggest that at least 700,000 people die annually from drug-resistant infections and, in the absence of measures, this number could rise up to 10 million by 2050, by far surpassing cancer as the major cause of death worldwide (Collaborators, 2022). In particular, infections caused by the ESKAPE pathogens, namely, *Enterococcus faecium, Staphylococcus aureus, Klebsiella pneumoniae, Acinetobacter baumannii, Pseudomonas aeruginosa* and *Enterobacter* spp., are increasingly associated with therapeutic dead-ends. As such, they constitute the top World Health Organization’s priority list of bacteria for which novel therapeutics are urgently needed. Identification and characterization of antibiotics with novel molecular scaffolds, innovative targets and/or original mechanisms of action, associated with a low potential of resistance induction, are an emergency, especially towards Gram-negative pathogens. Indeed, in the last decades, only six new classes of antibiotics have been approved (Butler et al., 2017). Recent efforts have begun to reactivate antibiotics research, and compounds are explored either through drug repositioning or through molecular mechanisms that are usually redundant with those of traditional antibiotics. This is the case for finafloxacin, a fluoroquinolone antibiotic that was recently approved to treat ear infections with *P. aeruginosa* (McKeage, 2015)(Randall et al., 2016), darobactin, which specifically targets Gram-negative bacteria (Imai et al., 2019), teixobactin, which is only functional against Gram-positive bacteria (Ling et al., 2015), or the new compound irresistin-16, which shows bactericidal activity towards both Gram-negative and Gram-positive bacteria without inducing any resistance, at least so far (Martin et al., 2020). These data pave the way for the discovery of new molecules, a mandatory step to provide efficient antibiotics acting on novel targets and thereby avoiding any cross resistances.

In this context, this study focuses on an innovative bacterial target, the Mutation Frequency Decline protein (Mfd), and its inhibition by novel small molecules. Mfd is a non-essential transcription repair coupling factor, ubiquitous in bacteria and absent in eukaryotes (Deaconescu, 2021). Mfd recognizes the RNA polymerase (RNAP) stalled at non-coding lesions and utilizes ATP to translocate along DNA, most likely forcing RNAP forward and ultimately dissociating it from the bulky DNA template (Roberts and Park, 2004). Mfd then stimulates DNA repair by recruiting components of the Nucleic Excision Repair system (Deaconescu et al., 2006; Million-Weaver et al., 2015; Pomerantz and O’Donnell, 2010; Smith and Savery, 2008). The structure of Mfd has been solved revealing the presence of an ATP-ase motor module, which is highly conserved among bacteria and functionally mandatory (Brugger et al., 2020; Deaconescu et al., 2006). Stress-induced mutagenesis can assist pathogens in generating drug-resistant cells during antibiotic therapy. Mfd has been associated with the development of antibiotic resistance in *Campylobacter jejuni* and *Helicobacter pylori* (Han et al., 2008; Lee et al., 2009) and suggested as an evolvability factor that promotes hypermutation in bacteria, thus accelerating the evolution of antimicrobial resistance (AMR) (Ragheb et al., 2019). These data strongly suggest that blocking Mfd could hamper molecular evolution, hence inhibiting resistance development in many bacterial pathogens.

Furthermore, Mfd is critical for virulence in the Gram-positive *Bacillus cereus* and in the Gram-negative *Shigella flexneri* species and confers resistance to nitric oxide (NO) stress (Darrigo et al., 2016; Guillemet et al., 2016). Production of reactive nitrogen species is a key step in the immune response following infections (Porrini et al., 2020). NO induces lesions in the bacterial DNA, thus limiting bacterial growth within hosts. Mfd is involved in the DNA repair following DNA damages and is thus required for bacterial resistance to the host response. Consequently, in both species, a mutant lacking Mfd is severely impaired in its virulence capacity in cell and animal models. As Mfd is widely conserved in the bacterial kingdom, these data highlight a mechanism – possibly ubiquitous in bacteria – to overcome the host immune response and tackle its mutagenic properties.

The function of Mfd in the bacterial bypass of host immunity, its role as an evolvability factor during antimicrobial resistance, its ubiquity in the prokaryotic world and its absence in higher eukaryotes, all support the selection of this protein as an innovative bacterial target for anti-infective therapies. Molecules that inhibit Mfd activity would neither directly inhibit nor kill bacteria, but instead curb bacterial evolution while impeding their ability to resist to the host immune response. Thereby, Mfd inhibitory compounds should boost the immune system response against pathogenic bacteria, while acting exclusively within the inflammation site, thus reducing collateral damages.

Here we report a targeted and structure-based high throughput *in silico* screening on the druggable ATP-binding site of Mfd to identify molecules capable to act as competitors of the cognate ligand ATP, thus inhibiting Mfd activity. One molecule, NM102, was identified as a potent molecule that specifically targets the ATP binding site of Mfd. *In silico* molecular analysis and *in vivo* assays revealed that NM102 displays antimicrobial activity, specifically in the context of infection, even against clinically relevant Gram-negative bacterial pathogens such as resistant *K. pneumonia*, *P. aeruginosa* and pathogenic *E. coli*. In animal models, NM102 blocked infections with these pathogens, without inducing toxicity to the host. Finally, NM102 blocked the function of Mfd as an evolvability factor, thus reducing the bacterial capacity to induce antimicrobial resistance.

As a whole, our findings identify and characterize a promising antimicrobial candidate that could be used either alone as a helper to the host immune system or in combination with other antibiotics by inhibiting Mfd evolvability activity.

## Results

### Identification of promising hits targeting Mfd in silico

In order to identify antimicrobials with a novel mechanism of action, we performed a rational, targeted and structure-based high throughput *in silico* screening of small molecules to challenge their efficiency to accommodate to the ATP binding site of the bacterial protein Mfd. A 3D modeling of Mfd of *E. coli* in an active conformation was designed (Supp Fig.1). Using this template, a large library of 4.8 million compounds was screened for their capacity to virtually bind the ATP-ase site of Mfd. The virtual screening led to the selection of 95 molecules for subsequent experimental validation (Supp Fig.1).

### Inhibition of Mfd activity in vitro

The 95 molecules were challenged to inhibit the ATP-ase function of *E. coli* Mfd *in vitro* (Fig. 1A). The inhibition rates of Mfd activity ranged from 10% to 85%. The best hit, called NM102, showed an inhibition rate above 80% and was therefore selected for further analysis. Dose-response assays on ATP-ase activity (Fig. 1B) and Lineweaver Burk plot (Fig. 1C) revealed a competitive mode of inhibition of NM102 to ATP with an IC_50_ of 29 µM and a K_i_ of 27 µM.

**Fig 1-.**
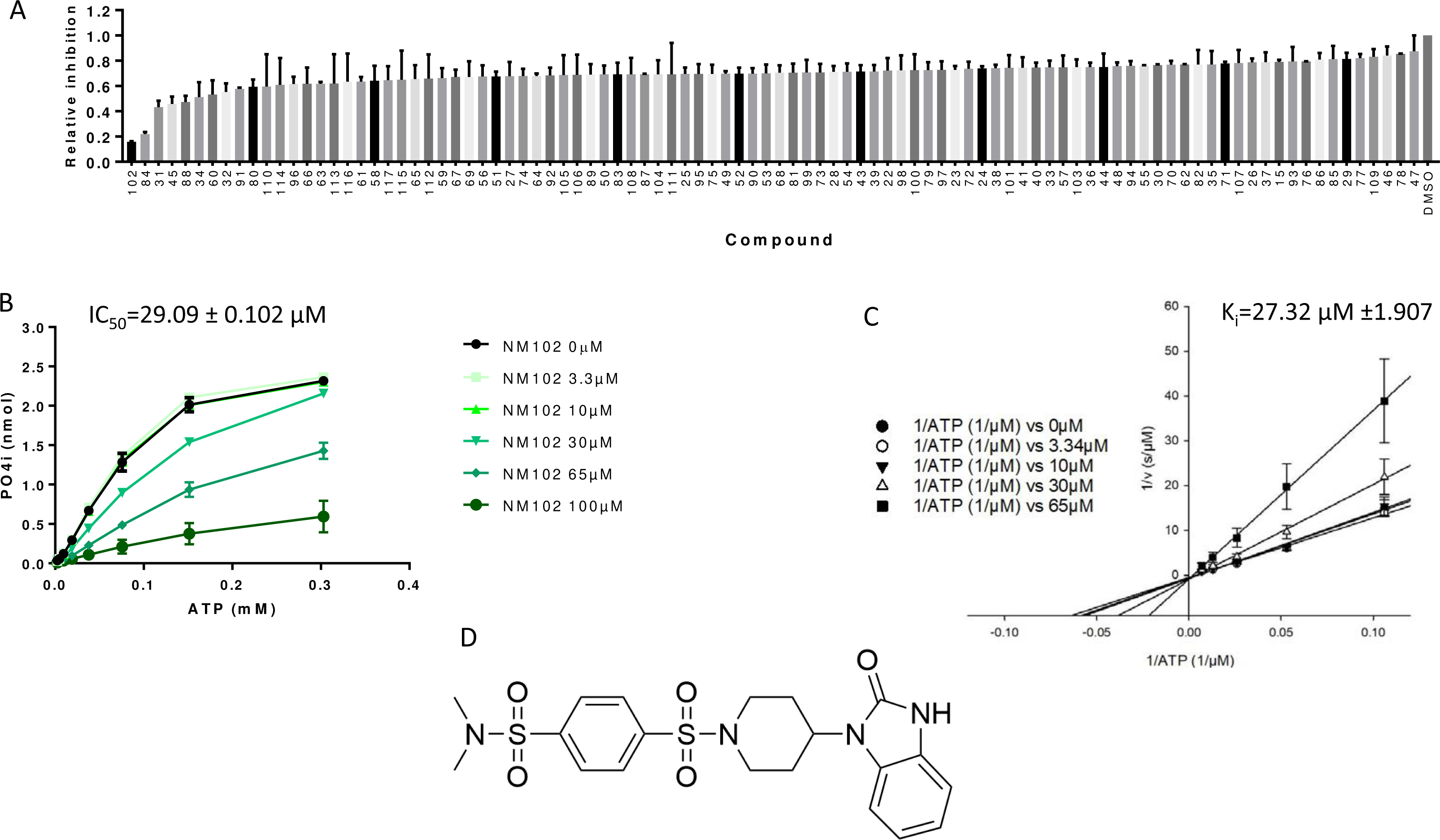
NM102 identification as ATP competitor for Mfd activity. (A) *In vitro* high throughput screening for inhibitors of Mfd-C ATP-ase activity. The indicated compounds were tested at a final concentration of 100 mg/mL. The data were normalized to those of the DMSO control. MfdC ATP-ase activity was measured at 0.35 µM at an ATP concentration of 1 mM. The results are the average of two independent experiments done in duplicate with standard deviation. (B) Michaelis Menten plot for inhibitory activity of NM102 (0 to 100 µM) on *E. coli* Mfd-C (0.35 µM) ATP-ase activity with ATP (0 to 0.3 mM). The results are the average of three independent experiments done in duplicate with standard deviation. The IC_50_ was computed using Graph Pad 7.05. (C) Lineweaver-Burk plot showing NM102 inhibition of Mfd-C ATP-ase activity through a competitive mode of action. (D) Chemical structure of the NM102 compound.

NM102 is a small molecule with a chemical scaffold relatively closed to ATP as it is composed of an indole-like ring close to the adenosine ring followed by a ribose-like ring and eventually polar sulfur groups that coul mimic phosphate moities (Fig 1D).

To assess the specificity of the NM102 inhibitor towards the ATP-ase activity of Mfd, we evaluated its capacity to decrease the ATP-ase activity in the eukaryotic ATP-ase yUpf1 protein (Fig 2A). NM102 sharply inhibited Mfd ATP-ase activity, while it showed no effect on yUpf1 activity.

**Fig 2.**
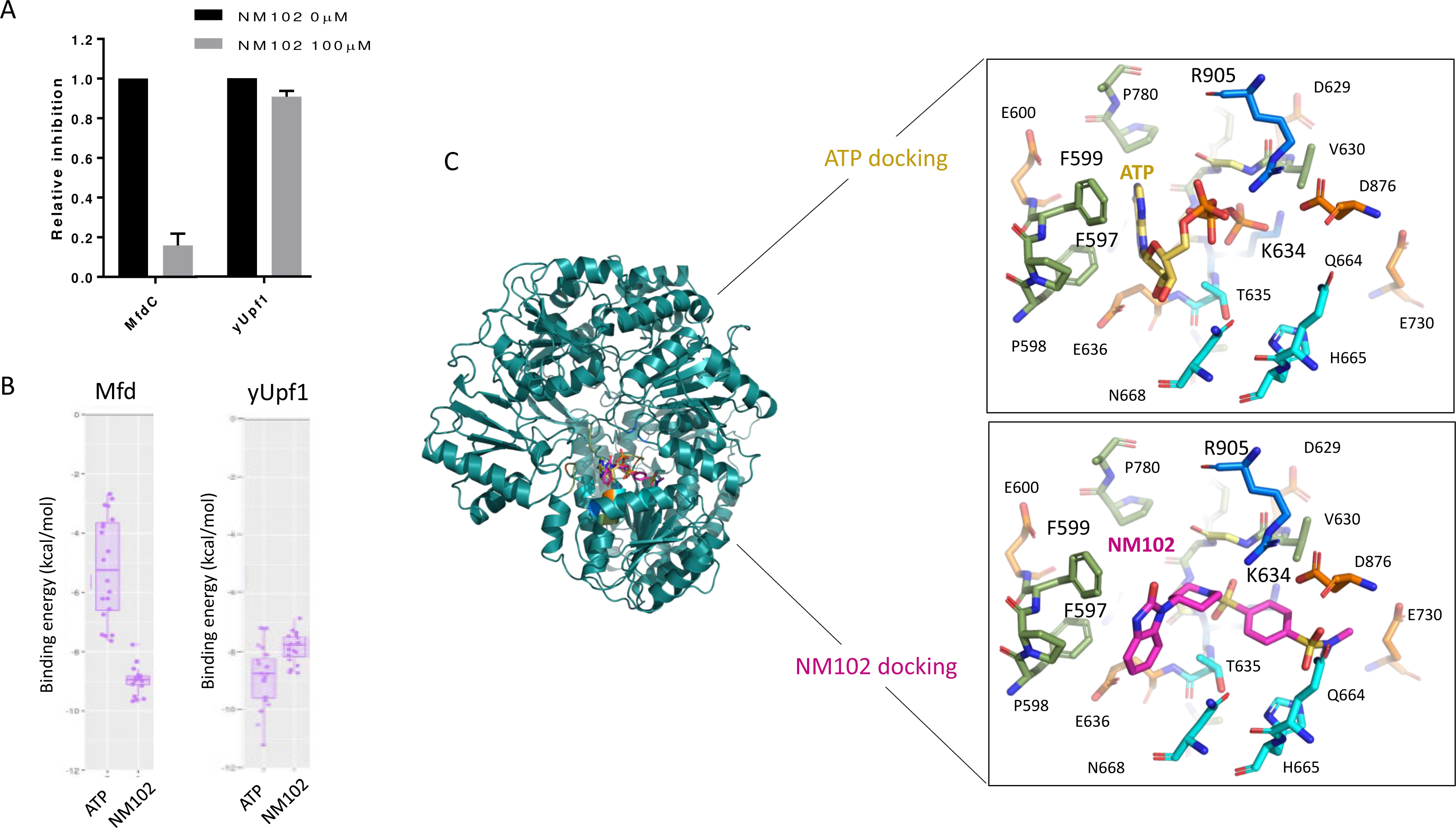
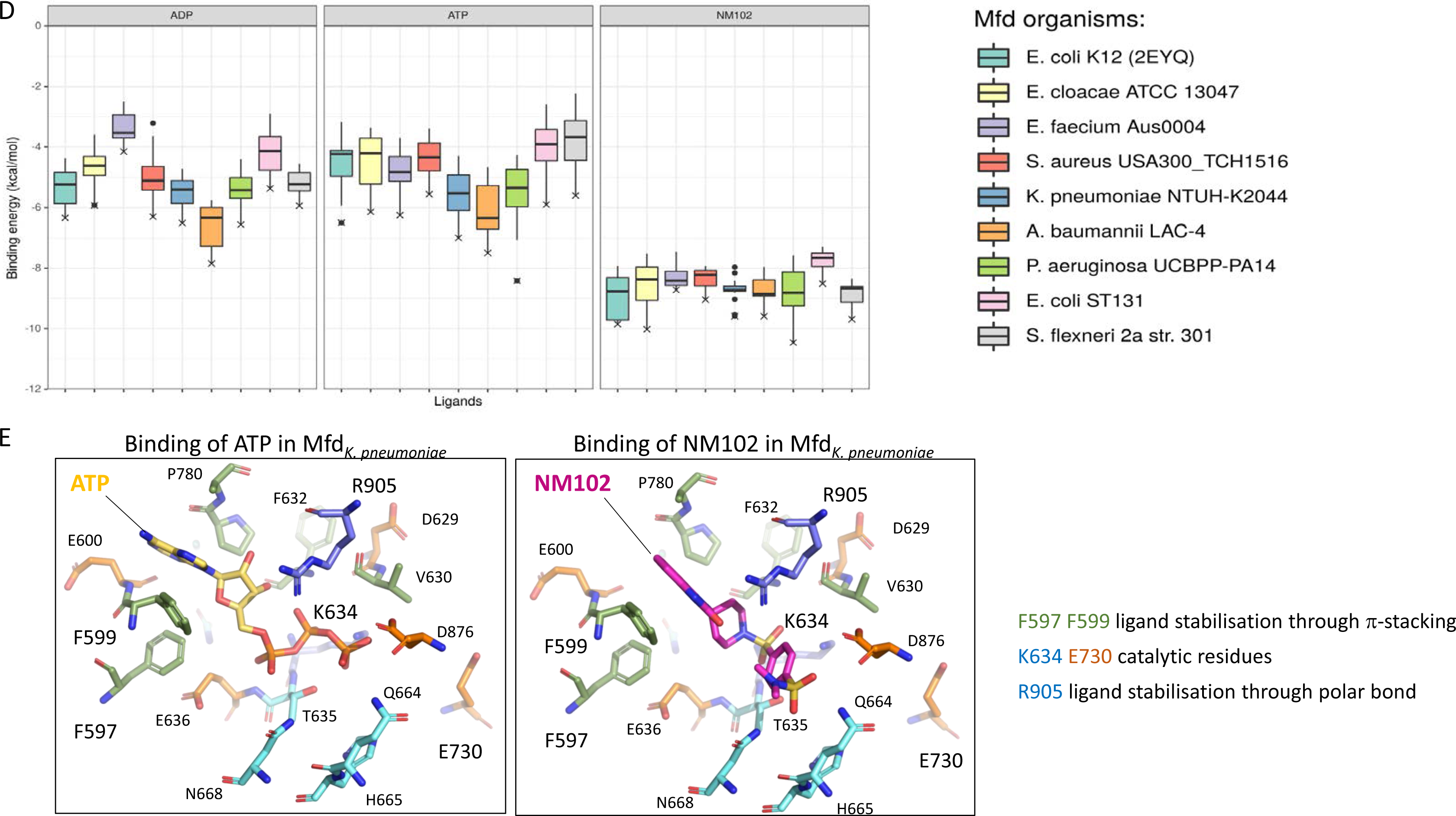
NM102 specifically binds the ATP binding site of Mfd. (A) NM102 inhibition of bacterial Mfd and eukaryotic yUpf1 ATP-ase activity. ATP-ase activity of Mfd and yUpf1 in the absence and presence of 100 µM of NM102 as assessed by using BIOMOL® Green reagent microtiter-plate assay. Data were normalized to that of the DMSO control. The graph shows the mean ± SD of three independent experiments. (B) Binding energy measured *in silico* in kcal/mol between Mfd and ATP or NM102 (left), and between yUpf1 and ATP or NM102 (right). (C) Left side: 3D structure of modelized full-length Mfd from *E. coli* in complex with NM102 and ATP at the active site. Right side: close view of ATP (upper panel, yellow stick) and NM102 (lower panel, pink stick) poses. The positions identified as conserved and involved in the binding of ligands are indicated: residues hydrophobic, acidic, polar, basic and glycine are colored in green, salmon, cyan, wheat and marine, respectively. (D) Binding energy measured *in silico* in kcal/mol between Mfd and ADP (left), ATP (middle) and NM102 (right) for the homology modeled Mfds from ESKAPE bacteria. (E) Close-view of ATP (yellow stick) and NM102 (pink stick) docked in Mfd of *K. pneumoniae*.

### NM102 binds Mfd of E. coli in silico with a better affinity as compared to ATP

In parallel with the experimental evaluation of the specificity of NM102, docking was computed in the ATP pocket of Mfd of *E. coli* and the binding affinity was evaluated (Fig 2B). NM102 showed a marked enhanced affinity for Mfd as compared to its cognate ATP ligand, with a binding energy of −9.8 kcal.mol^−1^ *vs* −6.5 kcal.mol^−1^, respectively. The docking highlighted that Mfd accommodates NM102 in its ATP binding site, within the same binding pocket encompassing most of the walker A motif (residues from Asp629 to Glu636), especially including the catalytic Lys634 and involving similar residues such as Phe597 and Phe599 through conserved π-stacking interactions, and Thr635 as well as Glu636 plus Arg905 for polar and charged interactions, respectively (Fig 2C). Markedly, we were able to define 21 conserved positions of close interactions involved in the binding of ATP and NM102 (Supp Table 1). Because NM102 is a slightly longer molecule as compared to ATP, its accommodation extends towards residues that are remote from the very catalytic nexus, namely Gln664, His665, Asn668 and even Glu730 of walker B motif, and D876 of motif V. This feature could explain the computed increase in affinity for NM102 as compared to ATP.

Furthermore, to confirm this specificity, NM102 was docked in the ATP pocket of the yUpf1 protein of *S. cerevisiae* and the binding affinity was measured (Fig 2B). In contrast to Mfd binding, NM102 shows a lower binding affinity to the eukaryotic yUpf1 protein as compared to the ATP control (−8.7 *vs* −11.2 kcal.mol^−1^ respectively), further suggesting the specificity of NM102 towards Mfd, which is consistent with our experimental data.

### In silico, NM102 binds to the Mfd active site of the ESKAPE bacteria

Mfd is widely conserved in the bacterial kingdom (Roberts and Park, 2004). To assess whether NM102 could more generally bind and inhibit the function of Mfd from a wide range of bacteria implicated in AMR, multiple sequence alignment and homology modeling of Mfd were performed for one strain of each species of the ESKAPE bacteria (Supp Fig 2 and 3). The alignment revealed that Mfds cluster distinctly into Gram-positive and Gram-negative bacteria but overall share 38% of sequence similarity along 1200 amino acid residues of the ESKAPE bacteria. Markedly, this average on Mfd full length sequence does not reflect the trend observed for each functional module. Indeed, modules composed of domains D1a-D2-D1b, D3, D4, D5-D6 and D7 show 21%, 14%, 47%, 66% and 26% of sequence similarity, respectively. Particularly, the ATP-ase motor module, composed of D5 and D6 domains, is by far the most conserved. It shares 66% and 36% sequence similarity and identity within the ESKAPE strains, respectively. Furthermore, the ATP binding site, which locates at the interface between D5 and D6, shows fairly high to strict conservation of the catalytic motifs, the so-called I to VI patterns as well as walker A & B boxes (black squared in Supp Fig 2).

**Fig 3.**
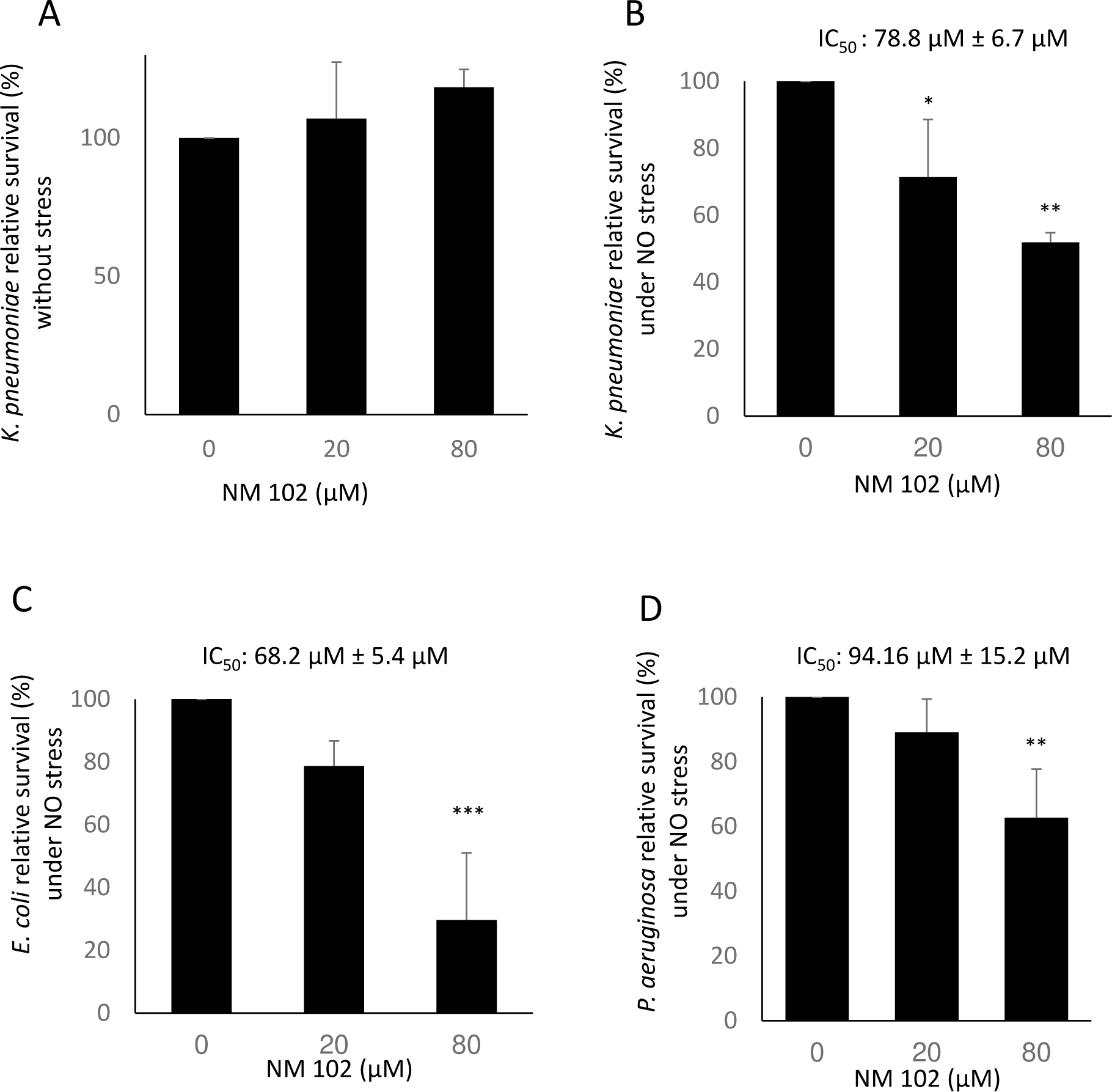
Antimicrobial effect of NM102 on Gram-negative bacteria during NO-stress. Strains were grown to exponential phase in LB medium. Bacteria solution was prepared in RPMI medium and dispatched in 96-wells plate. Bacteria were exposed for 4 h at 37°C to 50 µL of increasing concentrations of NM102 alone (A: *K. pneumonia* DOU) or with NOC5 as a NO donor (B: *K. pneumonia* DOU, C: *E. coli* ATCC 25922, D: *P. aeruginosa* CIP27853). Bacteria survival rate was calculated by normalizing bacteria load against control without NM102. The results reported are mean ± SD of four independent experiments each in triplicates, P values are calculated against the condition without NM102, using One Way ANOVA (**** P<0.0001, *** P<0.001, ** P<0.01, * P<0.05). IC_50_ were computed using Graph Pad 7.05.

To profile the affinity of NM102 towards Mfds from the ESKAPE bacteria and identify the probable conserved features of binding, NM102 was docked into each 3D homology model Mfd and its binding energy was measured (Fig 2D, E, Supp Fig 3). Molecular binding revealed that both ATP and NM102 engage in conserved interactions in the ATP binding site for all the modeled Mfds. Indeed, 21 residues were identified as positions always involved in the coordination sphere. All are strictly conserved except P2, P3, P9 and P19 which are nevertheless higly conserved (Suppl Table 1). The positions describe an hydrophobic pattern of interactions with Phe597 and Phe599 located around the adenine moiety, polar interaction with Thr634 and charged interaction with Glu636 and R905 around the phosphate groups of the ATP, respectively. The ATP binding ranges from −5.5 to −8.4 kcal.mol^−1^, whilst NM102 binding ranges from −8.5 to −10.5 kcal.mol^−1^, suggesting that NM102 binds to all Mfd from the ESKAPE group with a significantly better affinity as compared to the physiological ATP. Noticeably, this could indicate a putative universal inhibition power for NM102 towards this ATP-ase motor. Taken together, our results strongly suggest that NM102 targets the ATP binding site of Mfd protein of all ESKAPE bacteria, competing with physiological ATP, hence specifically inhibiting Mfd ATP-ase activity. As the challenging urgency is to identify molecules efficient against Gram-negative pathogenic bacteria of the ESKAPE group, the rest of the study is focused on clinically relevant *K. pneumoniae*, *P. aeruginosa* and *E. coli* strains.

### NM102 has antimicrobial activity in the context of nitric oxide stress

Mfd is a non-essential protein in unstressed conditions. However, we have previously shown that Mfd is essential to resist NO stress during infection (Darrigo et al., 2016; Guillemet et al., 2016). Our innovative antimicrobial strategy is to identify inhibitors with no antibiotic activity in unstressed conditions and active only in stressed conditions when Mfd is required. NM102 was thus first tested for its antimicrobial activity in the absence of stress. Under these conditions, no antimicrobial activity on *K. pneumoniae* was observed (Fig 3A). Conversely, NM102 inhibited clinical *K. pneumoniae* survival under NO stress conditions (IC_50_ of 78.8 µM) (Fig 3B). These results highlight that NM102 has an antimicrobial activity against *K. pneumoniae* under NO stress conditions.

When further tested towards other important bacterial pathogens, NM102 inhibited bacterial survival of clinical *E. coli* under NO stress conditions compared to unstressed conditions (IC_50_ of 61.2 µM, Fig 3C) and, although to a lesser extent, of *P. aeruginosa* (IC_50_ of 94.2 µM, Fig 3D). These data show that NM102 acts against Gram-negative pathogenic bacteria under conditions that mimic immune stress. Overall, NM102 proved to be an antimicrobial molecule active against clinically-relevant pathogens, including multidrug-resistant bacteria that may be associated with difficult to treat infections and sometimes to therapeutic failure.

### *NM102* is effective *in vivo*

To assess the activity of NM102 as an antimicrobial *in vivo*, *Bombyx eri* larvae were injected with *P. aeruginosa* CIP27853 and treated with NM102 (Fig 4A). The insects were then crushed and Colony-Forming Units (CFU) were counted by plating. Treatment with NM102 led to a drastic shift in the bacterial load showing that NM102 is effective against *P. aeruginosa in vivo*.

**Fig 4.**
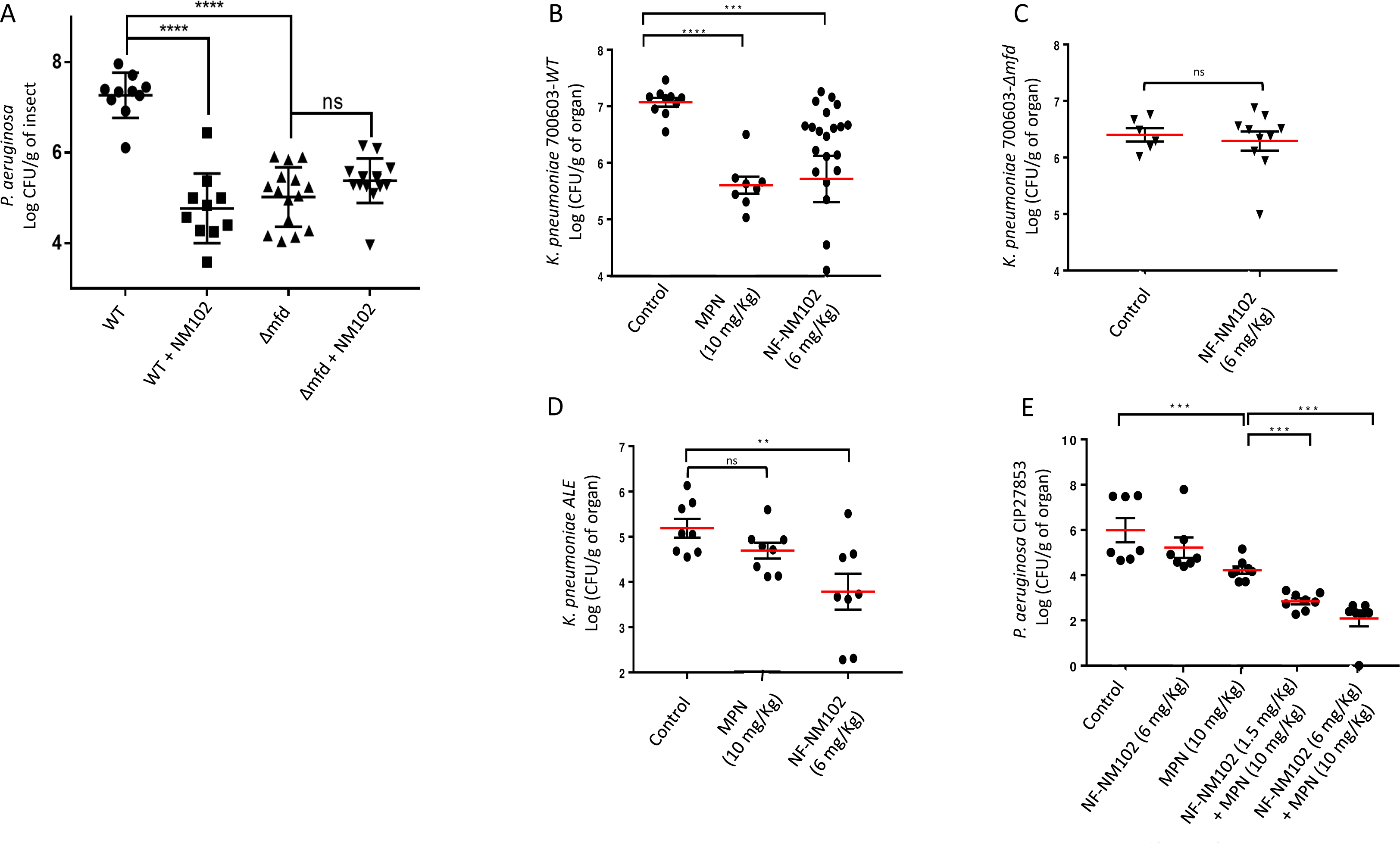
Antimicrobial effect of NM102 *in vivo* in insects and mice. (A) *P. aeruginosa* CIP27853 wt and Δ*mfd* strains were grown to late exponential phase in LB medium at 37°C and injected into forth instar silkworm larvae without or with NM102 (2.78 µg/larvae). Boosts of NM102 (6.97 µg/larvae) were administrated at 4 h and 8 h post infection. At 24 h post infection larvae were crushed and the content of each tube was serially diluted on LB plates for CFU numeration. Each point represents one larva. P values are calculated against the condition wild type-strain without NM102, using Mann-Whitney test (**** p < 0,0001). For mice experiments, *K. pneumoniae* ATCC 700603 wt (B) and Δ*mfd* (C), *K. pneumoniae* ALE (D), *P. aeruginosa* CIP27853 (E) were grown to late exponential phase and diluted in PBS. Intranasal (i.n.) bacterial administration was performed through slow instillation of 20 µL of bacterial suspension (1x 10^7^ CFU). 200 µL of NM102 formulation (NF-NM102, 6 mg/kg) or empty formulation (control) were administrated in mice via the i.p. route (B, C). Alternatively, 20 µl of a mixture containing bacterial suspension (1x 10^7^ CFU) and NM102 formulation (1.5 or 6 mg/kg) or empty formulation was administred via the i.n. route (D, E). MPN (10 mg/kg) was inoculated via the i.p. route. Mice were sacrified by cervical dislocation after 24 h and the log CFU in the lung was calculated per gram of organ. Each graph represents a pool of 2 to 4 independent experiments. Each dot represents a mouse. P values are calculated using Mann-Whitney test (*** p < 0,0005; ** p < 0, 001; * p < 0, 005).

To confirm that NM102 targets Mfd *in vivo*, the effect of NM102 was investigated on the survival of the *P. aeruginosa* mutant *Δmfd*. First, the amount of CFU of the *Δmfd* mutant was lower compared to the CFU recovered following insect infection with the wild-type strain, showing that Mfd is a critical virulence factor for *P. aeruginosa.* Second, NM102 did not decrease the bacterial load of the Δ*mfd* mutant, confirming that NM102 is specifically active on Mfd *in vivo*.

### NM102 nanoformulation is a potent antimicrobial agent active in vivo against clinically-relevant pathogens

To further investigate the potential of NM102 *in vivo*, a mouse lung model of infection with *K. pneumoniae* and *P. aeruginosa* was chosen (Liu et al., 2019). Due to its poor solubility in most common solvents (Supp Table 3), the above studies were performed using a solution of NM102 in DMSO. However, DMSO is known to be toxic to mice at high concentrations (Yi et al., 2017). Thus, an alternative to the DMSO vehicle was developed to further assess the therapeutic effect of NM102. Herein, a DMSO-free nanoformulation was prepared *via* a sonication assisted-nanoprecipitation method in the presence of poly(D-L lactide-*co*-glycolide) (PLGA) (Supp Fig 4A-D). The resulting NM102 nanoformulation (drug content = 0.6 mg/mL) exhibited an average diameter = 180 nm, with narrow particle size distribution (polydispersity index, PDI = 0.2) and a neutral surface charge (ζ potential = −2 ± 1 mV). An empty nanoformulation was prepared according to the same protocol but without the drug (average diameter = 136 nm ± 7, PDI = 0.08 ± 0.04, ζ potential = −4 ± 1 mV) and used as a control. NM102 nanoformulation showed colloidal stability for up to 2 weeks both at 4°C and at room temperature (Supp Fig 4E, F).

Mice were infected intranasally with *K. pneumoniae* and treated *via* the intraperitoneal route with the NM102 nanoformulation. After 24 h the CFU in the lung were assessed (Fig 4B). The NM102 nanoformulation significantly reduced the bacterial burden in lungs compared to control group treated with empty nanoformulation, showing that NM102 acts as an efficient antimicrobial against *K. pneumoniae* infection *in vivo*. The efficacy of the NM102 molecule was compared in the same conditions with that of meropenem (MPN), a beta-lactam antibiotic that is commonly used in the treatment of various infections including *K. pneumoniae* and *P. aeruginosa* (Liu et al., 2019). NM102 showed similar efficacy to MPN at 6 and 10 mg/kg in reducing the *K. pneumoniae* CFU in the lung of the infected mice (Fig 4B).

To confirm that NM102 targets Mfd *in vivo*, mice were also infected with the *K. pneumoniae* mutant *Δmfd*. The NM102 nanoformulation showed no effect on the bacterial burden of the Δ*mfd* mutant, further highlighting the inhibitor specificity (Fig 4C).

The efficacy of NM102 was further investigated on the clinical *K. pneumoniae* ALE strain, which is resistant to several antibiotics including MPN (Fig 4D).The NM102 nanoformulation at 6 mg/kg drastically decreased the bacterial load in the lung. Strikingly, MPN at 10 mg/kg had no significant effect on the resistant *K. pneumoniae* ALE, highlighting the potential of the NM102 molecule on resistant Gram-negative bacteria *in vivo*.

For *P. aeruginosa*, the NM102 formulation at 6 mg/kg had no significant impact on the bacterial load, whereas MPN at 10 mg/ml decreased the bacterial load (Fig 4E). The combination of the two treatments was more efficient against *P. aeruginosa* in an NM102-dose dependent manner compared to single treatments. These data suggest the possible use of NM102 as an antimicrobial drug during combination therapy.

### NM102 is not toxic for the host

In the development of antiinfectives, an important feature that needs to be assessed is their impact on the host. When tested for toxicity towards human cells (Fig 5A, B), NM102 and the NM102 nanoformulation did not induce cytotoxicity either to HeLa or to Vero cells at the highest doses used in insect and mouse experiments, respectively. MPN showed no toxicity to HeLa cells, but induced cytotoxicity in Vero cells at 10 and 100 µM.

**Fig 5.**
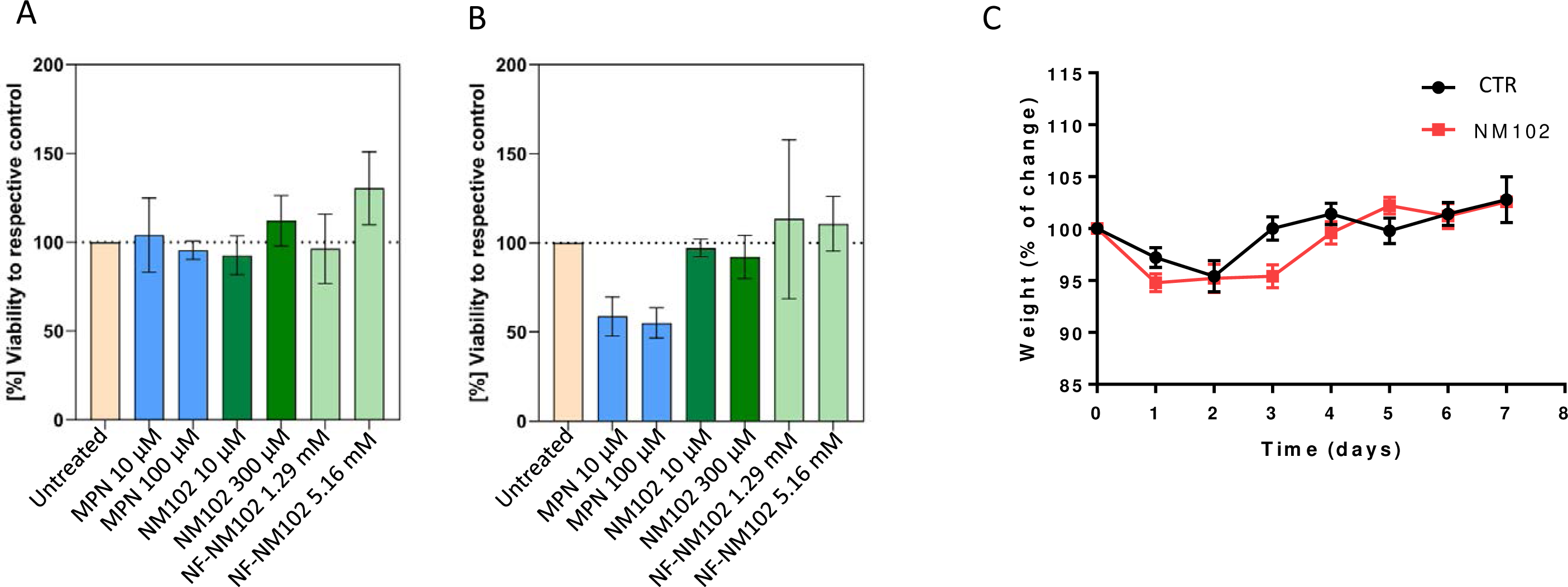
NM102 is not toxic to the host. HeLa (A) and Vero (B) cells were cultured in DMEM at 37°C and 5% CO_2_. Cells were treated for 1 h with MPN at 10 and 100 µM, NM102 at 10 and 300 µM, NM102 formulation at 1.29 and 5.16 mM or their respective control 10% DMSO or empty formulation. Cytotoxicity was assessed using CellTiter96®AQ_ueous_. Values for treated cells were normalized to the untreated control. The results reported are mean ± SD of three independent experiments each in triplicates (B) Mice were i.p. treated with NM102 (6 mg/kg) or control. Changes in the mice body weight were assessed for 7 days.

Treatment with NM102 had neither impact on insect or mouse survival nor on the mice weight (Fig 5C). Importantly, PLGA, which was used for the nanoformulation of NM102, is an FDA-approved copolymer for which neither systemic nor organ toxicity has been reported in animal models (Rong et al., 2014; Semete et al., 2010).

### NM102 may partially protect the host microbiome

As NM102 is active on bacteria only in the context of a stress like inflammation, we hypothesized that following pulmonary infection, the inflammation will be localized at the site of infection and that NM102 treatment should not affect the distant gut microbiota species, in contrast to treatment with conventional antibiotics.

To assess the innocuity of NM102 towards the gut microbiome *in vivo*, mice were first treated with NM102 without bacterial infection, and feces were recovered after treatment. The state of the microbiome was assessed by quantifying the overall quantity of bacteria in the feces by qPCR (Supp Fig 5). NM102 did not impact the overall quantity of bacteria in the feces, after 1 and 7 days of treatment.

We further assessed the composition of the gut microbiome following nasal infection by *K. pneumoniae* and NM102 formulation treatment. We did not detect changes in gut microbiome diversity between untreated and NM102-treated mice. The family-level microbiome profile, following NM102 treatment, largely resembled that of untreated mice (Fig 6A). The Shannon indexes of NF and NF-MN102-treated mice were similar to untreated mice (Fig 6B). Similarly, we did not detect significant differences in beta-diversity between treatments using weigted and unweigted UniFrac distances (Fig 6C). These results indicate that the nasal NM102 treatment does not induce changes in the richness or the structure of the gut microbiome.

**Fig 6.**
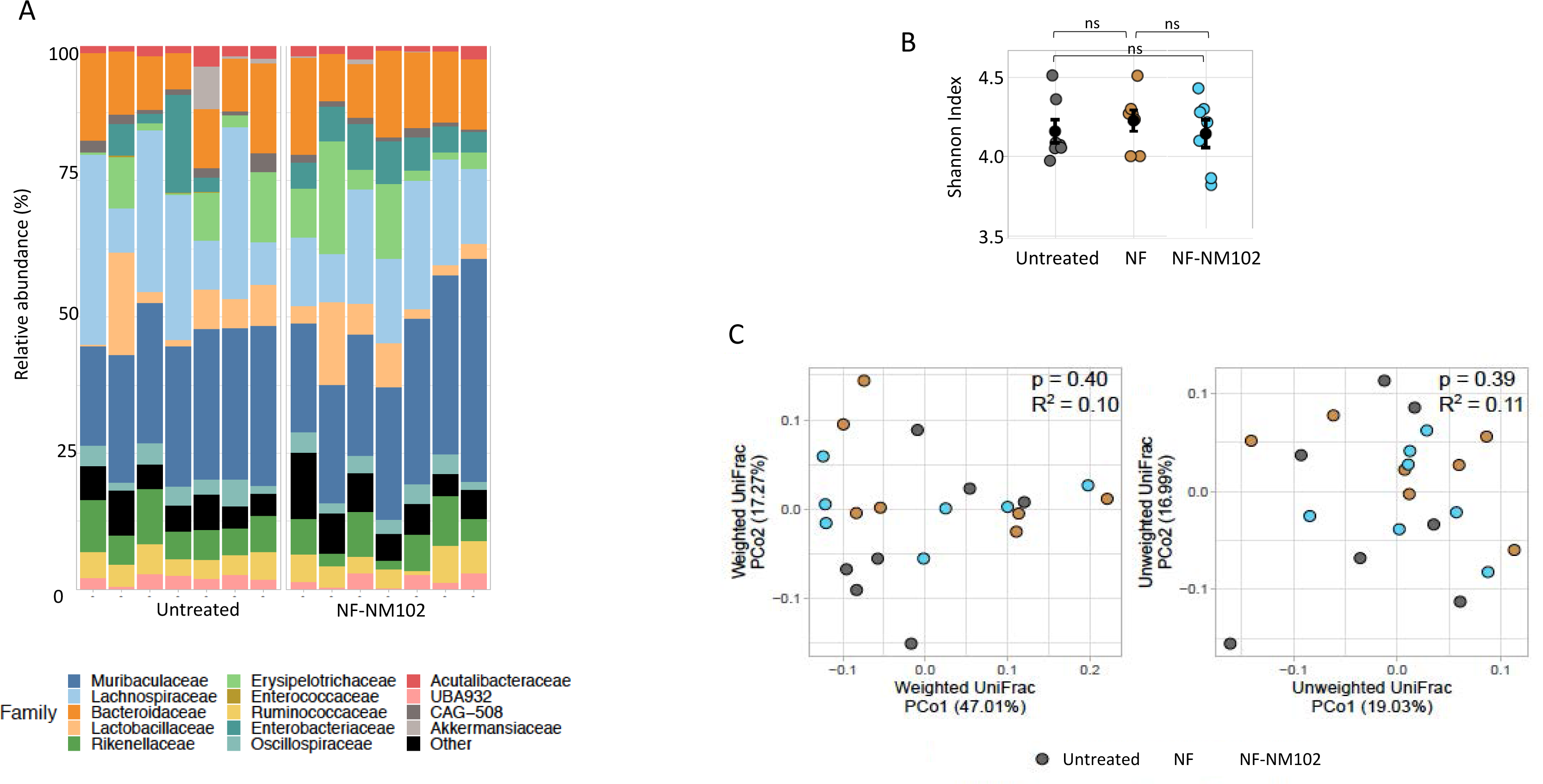
NM102 does not impact the microbiome. Gut microbiome diversity is unaffected by NM102. Mice were infected with *K. pneumoniae* and left untreated or treated with NM102 formulation (NF-NM102) or the empty formulation (NF) as control. After 24 h, the feces were collected and the diversity of the microbiome was analyzed by 16S rRNA sequencing. The relative abundance of the bacterial families (A), the Shannon index (B) and the profiling of the weighted and unweighted UniFrac distances (C) were determined. The bar colors represent bacterial families with mean abundance > 1%. Each dot (B, C) and each column (A) represents the value obtained for one mouse.

### NM102 is an anti-evolvability drug to combat AMR

Mfd has been shown to play a critical role in promoting bacterial resistance to commonly used antibiotics and has been proposed as a target for the development of anti-evolvability drugs (Ragheb et al., 2019). We thus assessed whether NM102 could also inhibit Mfd evolvability activity. First, the mutation rate of *E. coli* wild type and *Δmfd* mutant was assessed following exposure to Rifampicin (Fig 7A). The mutation frequency of the *Δmfd* mutant was significantly reduced compared to that of the wild type strain, confirming the role of Mfd in the induction of mutations. Second, the strains were treated with NM102 resulting in a drastic reduction in the mutation frequency of the wild type strain to a level similar to that of the Δ*mfd* strain. NM102 had no impact on the mutation frequency of the mutant strain.

**Fig 7.**
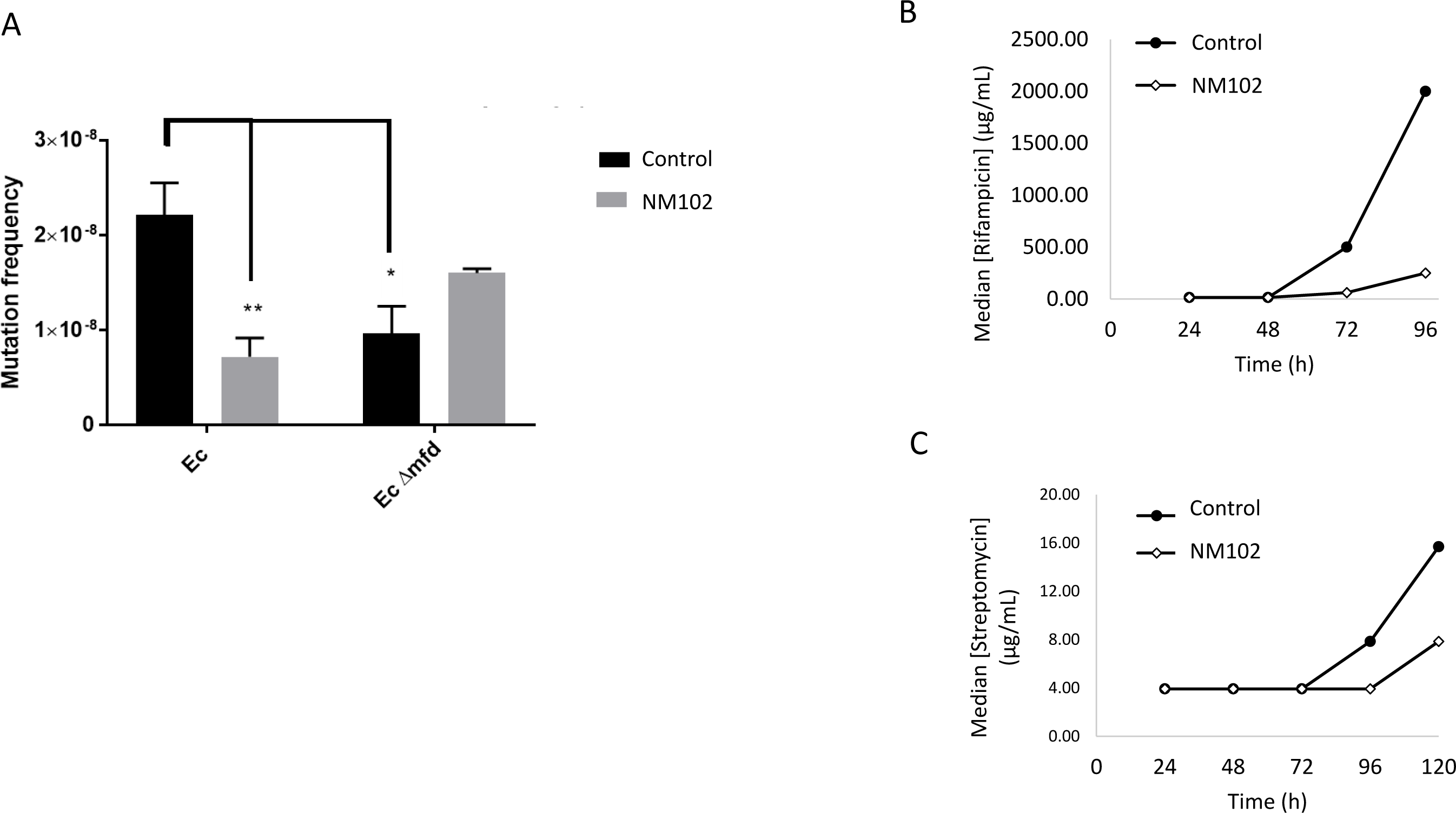
NM102 inhibits Mfd evolvability activity. (A) The mutation rate of *E. coli* wt and Δ*mfd* strains were measured following exposition to NM102 (100 µM) or the DMSO control, using the frequency of spontaneous accumulation of resistant mutants towards Rifampicin as measurement reference. Data in this figure are representative of at least three biological replicates. Error bars show SEM. Turkey’s multiple comparisons test was used to determine significance (∗p < 0.02; ∗∗p < 0.008). Resistance evolution of *E. coli* to Rifampicin (B) and Streptomycin (C) was measured following exposure to NM102 (100 µM) or the DMSO control. Line plots show median antibiotic concentration at each sampled time point for at least three independent experiments.

Then, the impact of NM102 on the evolution rate was evaluated following treatment of *E. coli* with several antibiotics. NM102 significantly decreased the evolution rate of *E. coli* following treatment with rifampicin (Fig 7B) or streptomycin (Fig 7C) by 8 and 2 folds, respectively. These data strongly suggest that NM102 targets the evolvability activity of Mfd and could thus be used as a combinatory drug to decrease antimicrobial resistance.

## Discussion

To address the need of novel bacterial drug targets and the design and synthesis of innovative anti-infectives with new mechanisms of action, especially towards Gram-negative bacteria, we have identified and characterized an innovative bacterial target, the Mutation Frequency Decline protein, Mfd, and promoted its inhibition by a novel therapeutic molecule. Mfd is a non-essential transcription repair coupling factor that is ubiquitous and conserved in bacteria but absent in eukaryotes. In this study, we identified a promising molecule, NM102, as an inhibitor of Mfd activity. We showed its dual capacity first to display antimicrobial activity in animals against *K. pneumoniae* and *P. aeruginosa*, two major nosocomial Gram-negative bacteria of the ESKAPE group, and second to inhibit Mfd evolvability function, thus reducing the frequency of antibiotic resistance appearance. As such, NM102 is active on its own to prevent bacterial infections. At the same time, it is also predicted to strongly reinforce the efficacy of existing antibiotics and prevent the emergence of resistance, an aspect rather unique and essential in the field of drug development and antibiotic therapy.

As Mfd is not an essential protein, standard MIC assays to identify inhibitory molecules could not be used. Thus, for the identification of the promising NM102 molecule, we developed a targeted approach based on the 3D structure of the active site of Mfd. This rational approach allowed to rapidly screen a large bank of molecules to identify potential hit inhibitory molecules. Hence, we propose a strategy that can be applied to other non-essential bacterial targets and provide a rapid and efficient pathway to screen small molecule potential without the development of laborious and specific *in vitro* activity tests. This structure-based method allowed us to predict the druggability of the target, based on the 3D structural descriptors (i.e. polarity, hydrophobicity, volume) of ligand-bound cavities in the Mfd ATP binding site (Rognan, 2017). The main advantage of such methods is their high interpretability in terms of pocket properties. In addition, once a potentially druggable pocket has been identified, it can be screened using various *in silico* tools to propose potential ligands for experimental validation and further optimization.

We chose to target the ATP binding domain of Mfd for the identification of inhibitory molecules because: i) ATP plays a central role in Mfd activity; ii) the 3D-structure of the ATP binding site structure in Mfd is known; iii) ATP-binding sites, despite their large structural diversity, are notoriously known to be druggable, as evidenced by inhibitors of many targets (protein kinases, HSP-90a for example) (Kahraman et al., 2007) and iv) although a large proportion of proteins carry ATP-binding sites, these sites present a large structural diversity and thus distinct specificity and selectivity towards ligands (Kahraman et al., 2007).

We report that NM102 is a potent ATP-like competitive inhibitor *in silico* and *in vitro*. *In silico*, its molecular mode of interaction grounds how NM102 is competitive and accommodates into the ATP pocket with a greater affinity than ATP. Markedly, its length could reinforce the bilobal binding between D5 and D6 domains, expected for ATP-ase activity, through direct and closer interaction with the Q664-H665 stretch and through ancillary salt bridge between the D876 and R905 residues. Consistently, NM102 competed *in vitro* with ATP. In addition, such interaction could trap Mfd in its ATP-ase active conformation and preclude any chance to engage into the ensuing DNA translocation step. Indeed, Mfd undergoes a functional cycle to couple ATP-ase to DNA translocation and then to switch towards UvrA recruitment. NM102 interacts strongly through polar interaction with the catalytic residue E730, of which mutation into a glutamine is defective in ATP hydrolysis. Besides, E730 is sandwiched between R733 and R953, and R953A has been reported to result in the loss of DNA translocation (Deaconescu, 2021). Any modification of interaction within E730 nexus, which connects both ATP-ase and DNA translocation activities, could impact the coordination of the two activities that must be exquisitely tuned. Markedly, the pseudo-adenine part of NM102 conserves a π-stacking with F597, similarly seen in the adenine moiety of ATP. Taken together, this work suggests that NM102 does not only occlude the ATP site but may also freeze the ATP-ase domain into a conformation that is unable to expel it, as it does for ADP, thereby blocking the functional cycle. These data provide mechanistic and molecular insights of NM102 binding with respect to its target and specificity.

Regarding specificity, NM102 neither binds nor inhibits the ATP-ase activity of the unrelated eukaryotic ATPase protein, yUpf1, further suggesting that the ATP pockets of ATP-ase proteins present structural diversity allowing distinct specificity towards ligands. The absence of toxicity of NM102, towards bacteria (in the absence of exogenous stress) and eukaryotic cells, provided further hints that NM102 did not affect general ATP-ase proteins activity.

Mfd activity enables the bacteria to overcome the host defense responses (Guillemet et al., 2016). The host defense against bacteria is predominantly mediated by cellular immune mechanisms. Various cells, such as macrophages, neutrophils and epithelial cells produce toxic species, including NO, during most types of infections. NO can severely damage biological molecules, such as proteins and nucleic acids, thereby inducing DNA damage and strand breaks (Akuta et al., 2006). NO is cytotoxic and mutagenic for various pathogens and host cells (Yoshitake et al., 2004; Zaki et al., 2005; Zhuang et al., 1998) and plays an important role during infections by limiting microbial proliferation (Porrini et al., 2020). Bacteria express sensor proteins that are able to detect NO and, accordingly, to switch on the expression of enzymes that detoxify NO before it reaches lethal levels (Porrini et al., 2021; Laver et al., 2010). In addition, we previously demonstrated a link between NO-induced bacterial DNA damage and DNA repair by Mfd (Darrigo et al., 2016). Mfd is ubiquitous in bacteria, and almost all bacteria induce a NO response from the host, so the Mfd-targeted inhibitor is expected to have a broad spectrum of activity. We have consistently shown that NM102 docks *in silico* to the Mfd of all ESKAPE pathogens, and that it has antibacterial activity in the context of NO stress on Gram-negative bacteria. As a proof of concept of efficacy against Gram-positive bacteria, we also confirmed that NM102 has antimicrobial activity against *B. cereus* enduring NO stress, the same *B. cereus* for which we had initially identified Mfd function during immune stress (Supp Fig 6).

Mfd activity is not restricted during NO burden. Indeed, it has been shown that Mfd protects against oxidative stress in *Bacillus subtilis* (Martin et al., 2019). This might explain the fact that NM102 was efficient *in vitro* during ATP test with quite low IC50 value, and also efficient *in vivo*, while providing higher IC50 values during NO *in vitro* stress, that may not fully represent the conditions encounterd by the bacteria during *in vivo* infections. The ability to evolve is critical for bacterial survival, especially in the context of pathogenesis, where escaping host immunity is essential and requires constant adaptation. Thus, Mfd may be involved in bacterial virulence, likely by preventing the DNA damages induced by the host through its overall immune response and/or stress conditions encountered *in vivo*. This question is difficult to technically address *in vivo* since bacteria encounter a lot of various stresses during infection. However, we could speculate that blocking Mfd activity by the inhibitory molecule hampers the bacterial capacity to cope with the host response (not only NO stress), allowing the immune system to be efficient against the invading pathogens.

We then focused our study on the impact of NM102 on the mutagenic property of Mfd to assess its role on antibioresistance. Indeed, Mfd has been demonstrated as a general evolvability factor promoting bacterial mutagenesis during infections (Deaconescu, 2021). It has been suggested that Mfd is required for developing high levels of drug resistance upon primary exposure to sub-inhibitory concentrations of antibiotics, accelerating AMR development (Ragheb et al., 2019) and that stress-induced mutagenesis can assist pathogens in generating drug-resistant cells during antibiotic therapy (Suzuki et al., 2018). Here we show that NM102 inhibits Mfd function as a mutation and evolvability factor during antibiotic stress. This decreases the ability of bacteria to develop resistance to classical antibiotics. As such, NM102 can be proposed as a new class of drug targeting the evolvability property of Mfd. By decreasing the resistance pressure, the drugs will have a longer efficacy, and this will in turn contribute to limit the spread of antibiotic resistance.

For clinical treatment, combination therapies have had and will still play an important role. We have shown that NM102 is efficient as antimicrobial *in vivo*, alone and during combination treatment with meropenem, and that NM102 treatment can kill bacteria in contexts where other therapies fail due to resistant strains. Thus, a combination treatment, including an inhibitor of Mfd coupled with a classical antibiotic, could constitute a promising strategy with an increased antimicrobial efficacy and combined with a reduced likelihood of resistance development at the onset of treatment. As recently proposed, targeting the evolutionary capacity of bacteria could also have wide-ranging implications outside of AMR development, from reducing cancer evolution to limiting pathogenic diversity in the context of host immunity (Ragheb et al., 2019).

Drug loaded into nanocarriers has been largely proposed for the delivery of poorly soluble drugs, allowing not only to overcome the physicochemical limitation of the drug, but also to ensure enhanced accumulation at the site of action and sustained drug release (Anselmo and Mitragotri, 2021; Hwang et al., 2020; Lee, 2020; Tibbitt et al., 2016). The design of a NM102 nanoformulation helped overcome the issues related to the low solubility of the drug molecule, allowing its administration in a clinically acceptable solvent. Herein, we provided a first proof of efficacy of the aqueous nanoformulation of NM102 opening the way to the development of a novel antibiotic formulation suitable for widespread use.

Classical antibiotics target essential bacterial pathways and as such, are highly efficient but not specific, thus also affecting commensal members of the microbiome (Maier et al., 2021). In particular, the gut microbiome is essential in human and animal health as it is involved in a variety of health associated processes, including digestion, metabolism, immune development, and also protects against invading pathogens. As Mfd is not an essential protein, its inhibition by NM102 should not have an impact on the distant host microbiome in the absence of inflammatory stress. To test this hypothesis, the gut microbiome composition was assessed after treatment with NM102 on healthy mice and on mice infected intranasally with *K. pneumoniae*. NM102 had no detectable effect on the diversity of the gut microbiome, which might be collaterally damaged during classical, non-selective antibiotic treatment. This is a rather unique feature and may help develop microbiome-friendly molecules that will not only heal the infectious diseases but also promote the general health of the patients. However, in the context of enteric infections, the local gut inflammation may render Mfd essential for the survival of bacterial commensals as well. Therefore, in those cases NM102-like compounds may also negatively impact the gut microbiome, imposing a collateral damage similar to that caused by the current antibiotics, with the advantage still of being effective in breaking resistance.

As a whole, new drugs targeting the non-essential, evolvability and virulence factor Mfd could be used alone or as supplemental drugs during the treatment of infections to improve the potency of current antibiotics and reduce the appearance of resistance. We are confident that these novel molecules could provide a new therapeutic option and expand the arsenal of drugs available to combat AMR and potentially other diseases.

## Ethic statement

All animals were handled in strict accordance with good animal practice as defined by the local animal welfare bodies (Unité IERP, INRAE Jouy en Josas, France, Agreement No. 78120) and all experiments were approved by the ethics committee COMETHEA and by the French Ministry of Higher Education and Research (APAFIS#10124-2017040413027917 v12). All animal experiments were performed in accordance with European directive 2010/63/EU. All efforts were undertaken to minimize animal suffering and to follow the 3R rules (Reduce, Refine, Reuse).

## Acknowledgments

We thank Antoine Allier for his help with R, Andrea Villarino for her input on the *in vitro* assay and Isabelle Barbosa for her input in the yUpf1 assay. We thank all members of the IERP animal facility (INRAE, Jouy en Josas) and Catherine Cailleau (Institut Galien, Châtenay-Malabry) for their technical help for animal studies. We thank Marion Leclerc for the qPCR experirments. We warmly thank Erwin Bohn and his team at the University of Tübingen for their help, advice and biological material for the construction of the *P. aeruginosa* Δ*mfd* mutant strain. We thank Cordula Gekeler for excellent technical assistance, Constance Porrini for her participation in the NO assay experimental design and Saoussen Oueslati for her help during the MIC tests. We thank the NCCT of Tübingen for sequencing. We warmly thank Dr. Libera Lo Presti for proofreading the manuscript.

This work was supported by *r*esearch grants from: SATT PARIS-SACLAY (CM2016-0067), Postdoctoral research Fellowship, Campus France (Prestige-2016-2-0011), Prematuration IDEX (CDE-2018-002324/IRE 2018-0022), DIM One health region Ile de France (R19104DD), Procope Campus France (46651WA), TWB biosciences and bioproduction, the Interdisciplinary Action "Health and Therapeutic Innovation" (HEALTHI) of the Université Paris-Saclay, the Mécénat des Mutuelles AXA and DFG (CMFI Cluster of Excellence EXC 2124).

## Material and Methods

### *E. coli* Mfd modelling

As we aimed to explore the capacity of Mfd to bind ligands possibly competitive to ATP, a prerequisite was to design Mfd in a conformation prone to accommodate a ligand, meaning that the ATP binding site must adopt an opened and accessible conformation. At the time of computation, all the available crystal coordinates of Mfd from *E. coli* were in an inactive form because of the so-called walker A motif that closed over the active site, hence precluding any binding (pdb id 2EYQ) (Deaconescu et al., 2006). The D2 domain of RecG helicase from *Thermotoga maritima* (pdb id 1GM5) is a structural homolog of Mfd, solved in active conformation complexed to ADP (Singleton et al., 2001). Thus, the nucleotide-bound active form of *E. coli* Mfd (K578-P780, UniProt identifier P30958) was obtained by morphing the inactive form of *E. coli* Mfd into the active form of RecG using standard settings of the Yale Morph2 server ().

### High-throughput *in silico* screening

The Bioinfo-DB database (http://bioinfo-pharma.u-strasbg.fr/bioinfo) is an in-house developed database of 4.8 million commercially available compounds as powder (1-50 mg) and filtered according to internal rules to contain drug-like compounds only. The database was first filtered to keep compounds grossly resembling to nucleotides and fulfilling the following properties: (i) at least one hydrogen-bond donor, (ii) at least two hydrogen-bond acceptors, (iii) at least one aromatic ring, (iv) predicted aqueous solubility higher than 50 µM (predicted with PipelinePilot v.9.5, Dassault Systèmes, Paris), (v) topological polar surface area lower than 120 Å^2^. The filtered set of 1.2 million compounds was then converted into three-dimensional (3D) space using Corina v3.40 (Molecular Networks, Erlangen, Germany). Up to 4 stereoisomers were created in case of the presence of undefined stereocenters. When the stereocenter was explicitly defined, it was kept unchanged. Altogether, 3D atomic coordinates were defined for 1 874 034 compounds, constituting the docking set.

The docking set was anchored to the above-described nucleotide-binding site of *E.coli* Mfd (Phe599, Thr602, Gln605, Gly631, Phe632, Gly633, Lys634, Thr635) with the Surflex-Dock v.3066 program (Jain, 2003). A prototyping molecular dynamics object or "protomol" was first generated from the list of binding site residues. Compounds were then docked with default settings (excepted for the -pgeom option) of the docking engine, keeping the best 10 poses according to the native Surflex-Dock scoring function.

Potential hits were selected according to the following two strategies (Supp Fig.1):

Strategy A: The 178 best scoring poses (Surflex-Dock score > 10) from the docking set were retained. Then, compound redundancy was removed (highest score retained if more than two poses originate from the same compounds) to yield 91 compounds. A chemical diversity selection by maximum common substructures was performed using the LibMCS algorithm (ChemAxon Ltd., Budapest, Hungary) to retain a first set (SET1) of 23 chemically diverse compounds.
Strategy B: All poses from the docking set were submitted to an interaction-based filter (Marcou and Rognan, 2007) to select 206 non-redundant compounds verifying absolutely the following interactions: hydrogen bond to Gln605.OE1 atom, hydrogen bond to Gln605.NE2 atom, hydrogen-bond to Gly631.N atom, π-π aromatic stacking to Phe599. In case of multiple poses from the same compound verifying the interaction similarity filter, the top scored posed (best Surflex-Dock score) was kept. A chemical diversity selection by maximum common substructures was performed using the LibMCS algorithm (ChemAxon Ltd., Budapest, Hungary) to retain a second set (SET2) of 74 chemically diverse compounds. Previously defined SET1 and SET2 were merged to yield a final selection of 95 unique hits (two hits were common to both sets) that were further purchased in 5 mg quantities.

### Homology modelling of Mfd from the ESKAPE species

Mfd sequences from the ESKAPE pathogens were retrieved from NCBI (https://www.ncbi.nlm.nih.gov/). They were aligned using the multiple sequence alignment (msa) MAFFT algorithm with default parameters (Katoh et al., 2002). The alignment visualization was done using Jalview 2 (Waterhouse et al., 2009).

For homology modelling, a chimeric Mfd was modeled to mimic full length Mfd from ESKAPE bacteria in a conformation compatible with ligand binding. During this study, the structure of Mfd from *Mycobacterium smegmatis* in complex with ADP (pdb id 6ACX) was solved in an active opened conformation. The structures of Mfd full length from *E. coli* in its apo form and Mfd full length from *M. smegmatis* in its holo form served as 3D templates. For the former, its ATP-ase center was excised and replaced by RecG helicase D2 domain to reinforce the coordinates of an active ATP-ase center. Then, homology modeling of the Mfds from the ESKAPE bacteria was performed by inserting their corresponding Mfd sequences into the building software Modeller (version 9.18) (Sali and Blundell, 1993) and by using our Mfd chimera as 3D template. For each Mfd, hundred models were computed and ranked according to the score function of Modeller. Eventually, the final one was selected upon this lowest score as it had the highest probability to satisfy the spatial contraints. Last, the stereochemistry of the model was checked using Procheck (Laskowski et al., 2018).

### NM102 docking

ATP and NM102 were docked individually into the active site of each modelled Mfd. AutoDock Tools 4.2 (ADT4) was used with a cubic grid box centered onto the active site of the ATP-ase, with the algorithm of Lamarck and the default parameters for 20 runs (22). ATP and NM102 coordinates were obtained from the ZINC database (https://zinc.docking.org/substances/home/) in mol2 format, converted into ADT4’s PDBQT format with their dihedrals set free to rotate. The crystal coordinates of the ADP molecule were first extracted from Mfd of *M. smegmatis* and redocked into this site, following the protocol described above. As the method docks it similarly to the coordinates of the solved structure, it serves as positive control, and subsequently the protocol was approved for docking of ATP and NM102 molecules in all homology modeled Mfds. ATP and NM102 ligands were also docked into the crystal structure of the eukaryotic protein yUpf1 (PDB ID: 2XZL) (Chakrabarti et al., 2011). For each run of each complex, an affinity score was calculated. Regarding this score, the most probable pose of ligand, that corresponds to the lowest interaction energy computed between protein and ligand, was selected. The binding energy was measured in kcal.mol^−1^. Finally, holo models were visually inspected using PyMOL 2.0.7. (Schrödinger, LLC) and protein-ligand interactions were characterized using the Protein-Ligand Interaction Profiler (PLIP) program (Adasme et al., 2021).

### Bacteria

*K. pneumoniae* DOU CTXM-15, ALE CTXM-15+, *P. aeruginosa* CIP27853, *E. coli* ATCC25922 were provided by Dr. Thierry Naas (CHU de Bicêtre, Le Kremlin-Bicêtre, France) (Supp Table 1). The minimum inhibitory concentration (MIC) of antibiotics was realized on those strains. Briefly, a bacterial solution of 0.5 McFarland was prepared in physiological water and then diluted 1/1000 in Mueller Hinton broth (MHB) medium. 50 µL of bacterial solution were added in 96 well plates containing 100 µL of various concentration ranges of antibiotics (Penicillin, Ampicillin, Ceftriaxone, Meropenem, Cefotaxin, Oxacillin, Streptomycin, Ciprofloxacin). The plates were incubated over night at 37°C. Bacterial growth was measured in each well to determine the MIC (Supp Table 2).

All Strains were stored at −80 °C as 20% (v/v) glycerol cultures.

### Construction of Δ*mfd* mutants

The *P. aeruginosa* CIP27853 Δ*mfd* mutant strain was constructed by double-crossover region deletion. Briefly, using the available sequencing information of *P. aeruginosa* CIP27853 strain, around 1-kb regions upstream (region 3360879 to 3361826) and downstream (region 3363535 to 3364310) of the targeted *mfd* gene were synthesized by the Genecust company (Boynes, France). The two fragments were cloned and juxtaposed into the (suicide) pEXTK vector by the Genecust company. The constructed plasmid was named pPa001 and transformed into electro competent *E. coli* SM10 λpir and then, by conjugation into *P. aeruginosa* CIP27853 strain as previously described (Hmelo et al., 2015). Briefly, the donor and receptor strains were mixed at a 1:2 ratio on a sterile membrane filter with a pore size of 0.05 μm (VMWP02500 Milipore) deposited on LB agar plates supplemented with irgasan 25 µg/ml and gentamicin 75 µg/ml for 18 h at 37°C. After mating, bacteria were resuspended from filter with LB-IPTG 1 mM, incubated for 3 h at 37°C and plated on LB agar plates with AZT 100 µg/ml and IPTG 1mM for 18 h at 37°C. The trans-conjugates were selected for loss of gentamicin resistance by picking individual colonies on LB agar plates and on LB supplemented with 75 μg/mL of gentamicin at 37°C. The deletion of the *mfd* gene by double recombination event was verified by PCR using primers located upstream (GAACACCACCAGTTCCACCTG) and downstream (GGTAGAGCTGGCCAATCAGC) of the cloned region and by sequencing. The corresponding mutant was named *P. aeruginosa* Δ*mfd*.

The *K. pneumoniae* ATCC 700603 Δ*mfd* mutant strain was constructed by double-crossover region deletion. Briefly, around 1-kb region upstream (region 2056347 to 2057137) and downstream (region 2060189 to 2060810) of the targeted *mfd* gene were synthesized using the available sequencing information of *K. pneumoniae* ATCC700603 strain, by the Genecust company (Boynes, France). The two fragments upstream and downstream of the *mfd* gene were cloned at each side of the chloramphenicol cassette (CatR) from the pDK3 plasmid (Addgene) into the pUC57 vector (Addgene) by the Genecust company. The synthesized DNA fragments and the CatR cassette were digested with the appropriate enzymes and assembled by ligation to produce a “*mfd*-upstream”-“CatR”-“*mfd*-downstream” *Kpn*I-*EcoR*I fragment, which was then inserted between the *Kpn*I and *EcoR*I sites of the pDG704 vector (Addgene). The constructed plasmid was named pKp002 and transformed into electro competent *E. coli* SM10 λpir and then by conjugation into *K. pneumoniae* ATCC700603 strain. Briefly, the donor strain and receptor strains were mixed at a 2:1 ratio on a sterile membrane filter with a pore size of 0.05 μm (VMWP02500Milipore) and deposited on LB agar plates for 4 h at 37°C. After mating, bacteria were resuspended from filter with 1.5 mL of LB medium. The trans-conjugates were selected on LB agar plates supplemented with 25 μg/mL of chloramphenicol at 37°C. The deletion of the *mfd* gene by double recombination event was verified by PCR using primers located upstream (CCCCAGGATGTTGTTTTGCA) and downstream (CCGTTATCCTGCACCACATG) of the cloned region and by sequencing. The corresponding mutant was named *K. pneumoniae* Δmfd.

### Chemicals

The 95 molecules were sourced from commercial vendors: Enamine, Chembridge, Chemspace, AbamaChem, Asinex, Chemdiv, Interbioscreen, BCH research and Specs. Each compound was dissolved in DMSO (Sigma-Aldrich) and stored at room temperature until further use. The Roswell Park Memorial Institute (RPMI) 1640 GlutaMAXTM medium was purchased from Gibco. The Nitric Oxide (NO) donor, 3-[2-hydroxy-1-(1-methylethyl)-2-nitrosohydrazino]-1-propanamine NOC-5, was purchased from Calbiochem, Sigma-Aldrich. NOC-5 was dissolved in NaOH 0.01 N at a final concentration of 200 mM. NOC-5 has a half-life time of NO release of 25 min at 37 °C. Under these conditions, a stable NO-amine complex can spontaneously release two NO equivalents (Porrini et al., 2020). Meropenem (MPN) was purchased from Sigma-Aldrich and dissolved in water at a stock concentration of 25 mM. Poly(lactide-co-glycolide) (PLGA) (Resomer® RG502 H, acid terminated, M_W_ = 7-17 kDa), poly(vinyl alcohol) (PVA) (87−89% hydrolysed, MW = 30−70 kDa), trehalose and all other reagents and solvents were supplied by Sigma-Aldrich (France).

### Protein purification

*E. coli* MG1655 His6-tagged MfdC (residues 451-1148 Uniprot accession code P30958) was synthesized and purified by BioBasic Inc. (Markham, ON, Canada) with a purity >90%. Briefly, expression plasmid PET21a containing *mfdC*, was transformed into *E.coli* BL21(DE3) and expression of His6-tagged MfdC was induced by 0.5 mM IPTG at 37℃ for 4 h. Cells were lysed and protein was purified with the Ni-IDA column. Elution fractions were dialyzed with dialysis buffer (2 mM DTT, 50 mM Tris, 300 mM NaCl, pH 8.0) overnight at 4℃.

*Saccharomyces cerevisiae* protein yUpf1 HD (residues 221–851 Uniprot accession code P30771, which corresponds to the helicase domain) was kindly provided by Dr Hervé Le Hir (Institut de biologie de l’Ecole normale supérieure (IBENS), Paris, France) and described in (Kanaan et al., 2018).

### Inhibition of ATP-ase activity *in vitro*

MfdC enzyme activity was evaluated by measuring the quantity of inorganic phosphate (PO4i) released using BIOMOL® Green reagent microtiter-plate assay (Enzo Life Sciences). For the initial drug screening, MfdC (0.35 µM) was incubated with DMSO (2% (v/v)) or with the 95 compounds (100 mg/mL) and 1 mM ATP for 10 min at 37⁰C. To further evaluate NM102 activity, MfdC (0.35 µM) was incubated with NM102 at the indicated concentration (0 to 100 µM). ATPase reaction was measured in Tris buffer 0,05M at pH 8, in the presence of ATP (0 µM to 0.3 mM) for 30 min at 37⁰C. yUpf1 HD ATPase activity was measured in reaction buffer (20 mM MES pH 6.0, 100 mM potassium acetate, 1 mM DTT, 0.1 mM EDTA, 1 mM magnesium acetate, 1 mM zinc sulfate, and 5% (v/v) glycerol). 0.127 µM of protein was incubated with 2% DMSO or 100 µM NM102. ATPase reaction was quantified after 30 min incubation at 30°C. For MfdC and yUpf1 HD, 50 µL of each reaction medium was transferred into clear, flat-bottom 96-well plates and the reaction was terminated by the addition of 100 µL of BIOMOL® Green reagent. The absorbance at 620 nm was measured in a microplate reader (Tecan). The absorbance values were then transformed into nmols of released PO4i based on a PO4i standard curve prepared as recommended by the supplier. The potency of each compound was calculated relative to the DMSO control.

The z’ factor of the ATP-ase test was calculated by using the formulae Z′=1−(3σc++3σc−)/(|μc+−μc−|) where (σc+) and (σc−) are the data standard deviation for the high and low reference control, respectively, and |μc+–μc−| is the absolute value of the difference of the two control signal means. Plot of MfdC ATPase activity versus ATP concentration (0.002-0.3 mM) in the presence of various concentrations of NM102 inhibitor was used to measure the IC_50_. Data analysis and IC_50_ computing were performed using the Graph Pad Prism 7.05 software (San Diego, CA). Lineweaver-Burk plots were obtained by using Sigma Plot Software.

### Nitric Oxid (NO) cell free assay

Bacteria were grown in LB medium at 37°C under agitation at 200 rpm until late exponential phase. Bacterial suspensions were prepared in RPMI medium at concentrations between 5 x 10^4^ and 1 x 10^5^ CFU/mL and 150 μL were dispatched into a 96-well plate. Bacteria were exposed to NO through the addition of 50 μL of NOC-5 at final concentrations ranging from 0 to 1000 μM. Bacterial survival rate was quantified by plating on LB agar after 4 h at 37°C and by normalizing bacterial load in NOC-5-treated samples against control untreated samples (0 μM of NOC5). NOC-5 concentrations (Log10-transformed) were plotted and a non-linear fit with a variable Hill slope was made using Graph Pad Prism 7.05 to obtain NO dose-response curves. NOC-5 concentrations required to induce 10 and 20% of mortality of the strain were calculated using R and the drc package (Christian Ritz and Jens C. Strebig).

For efficacy assays, bacterial solutions were prepared as described above. Bacteria were exposed to 50 µL of NOC5 at a concentration inducing a survival of around 80 - 90 % compared to the untreated condition (*K. pneumoniae* Kp DOU 14 µM, *P. aeruginosa* 5 µM, *E. coli* ATCC 42 µM), in absence (0 µM) or in the presence (from 5 to 80 µM) of NM102. Bacterial survival rate was quantified by plating on LB agar after 4 h at 37°C and by normalizing bacterial load in NM102-treated samples against control samples. Toxicity of the inhibitor in the absence of NO was assessed by exposure of bacterial suspension to NM102 at a concentration ranging from 0 to 80 µM of NM102 as described above, but without NOC5. NM102 concentration required to induce 50% of mortality of the strain (IC_50_) was calculated using R and the drc package.

### Nanoformulation of NM102 and characterization

NM102 was formulated in the presence of poly(D-L lactide-*co*-glycolide) (PLGA) according to a previously described method with some modifications (Van de Ven et al., 2012). It consists of an ultrasonic emulsification process, followed by dialysis to remove the organic solvent. First, 120 mg of PLGA and 28 mg of NM102 were dissolved in 4 mL of DMSO and then the organic phase was injected through a 23 G-needle into 50 mL of a 0.5% w/v aqueous solution of PVA under sonication in ice (5 min, amplitude 20%, cycle of sonication: 10 s ON and 10 s OFF) using a tip sonicator (Vibra-cellTM– 75041, Fisher Bioblock Scientific, Belgium). The organic phase was then removed by dialysis (24 h, room temperature) using a 100 kDa MW cut-off cellulose ester membrane (Thermo Fisher Scientific, France). Empty nanoformulation was prepared according to the same protocol but in absence of drug in the organic phase. Nanoformulations were stored at 4°C. Intensity-averaged diameters (*D*_z_) and particle size distributions were measured at 25°C by dynamic light scattering (DLS) using a Nano ZS instrument (173° scattering angle, Malvern, France) after 1/20 dilution in Milli-Q water (Millipore, France). The surface charge of the nanoformulations was determined by measuring the *ζ*-potential at 25°C using the same instrument after 1/10 NP dilution in 1 mM NaCl. All measurements were performed in triplicate. Colloidal stability of the nanoformulations was assessed by measuring the evolution of the mean diameter and the particle size distribution for up to two weeks at 4°C and room temperature.

For conservation, 20-mL flat bottom glass vials were filled with 2 mL of nanoformulation supplemented with 5% w/v of trehalose as a cryoprotectant. Samples were frozen at −20°C for 12 h and then freeze-dried (24 h) using an Alpha 2–4 LD plus freeze-drier (Martin Christ, Germany) (condenser temperature < 70°C, vacuum < 0.1 mbar). Freeze-dried samples were capped, weighted and stored at 4°C. Samples were reconstituted immediately before use by addition of Milli-Q water, followed by gentle homogenization by pipetting and vortexing.

The drug content was determined by direct quantification of NM102 in freeze-dried NM102 nanoformulation samples (without cryoprotectant). Briefly, one sample was dissolved in DMSO (1 mL) by vortexing. The amount of NM102 was then quantified by UV spectroscopy at 282 nm (Perkin-Elmer UV/VIS spectrophotometer, Germany). The drug concentration was calculated using a NM102 calibration curve in DMSO (correlation coefficient >0.999), as follows: drug content (mg/mL) = weight of drug/volume of nanoformulation dispersion.

### Insect assay

Fourth instar silkworm larvae *Bombyx eri* were purchased from *« L’office pour les insectes et leur environnement* » (OPIE), Guyancourt, France. Larvae were reared with *Ligustrum vulgare* or *Ligustrum japonicum* until they reached a weight of [0,6 – 0,9] g. Prior to assays, *Bombyx eri* larvae were kept under starvation overnight and randomly distributed in groups of 30. *P. aeruginosa* wt and Δmfd strains were grown in LB medium at 37°C under agitation at 180 rpm until late exponential growth phase. Bacterial cultures were then diluted in Phosphate Saline Buffer (PBS). The last dilution to obtain the desired concentration was done in PBS with 10% DMSO with or without NM102. Bacteria were plated right before infection to determine the quantity injected. Larvae were injected with 20 µL of bacterial suspension (3 x 10^3^ CFU/injection in 10% DMSO) containing NM102 (2.78 µg/larvae) or 10% DMSO in the haemolymph via the last pro-leg. Larvae were placed at 27°C. At 4 h and 8 h post first injection, larvae were injected with 50 µL of: (i) PBS, 10% DMSO or (ii) PBS, 10% DMSO with NM102 (6.97 µg/larvae) *via* the last pro-leg on the other side of the larvae. All injections were performed using a 1 mL hypodermic syringe with a 0.5 x 25 mm needle and an automated syringe pump (KD Scientific KDS 100). 24 h post infection, insects were placed in 2 mL sterile tubes containing 500 µL of PBS and 1.4 mm ceramic lysing beads (MP Biomedicals, Solon, OH, USA) and crushed using a Fast Prep instrument (MP Biomedicals, Solon, OH, USA). The bacterial content in each tube was assessed by serial dilutions on LB plates and CFU numeration. The log CFU was calculated per gram of larvae.

### Mice infections

Nine-week-old, specific-pathogen-free, female C56BL/6J mice were purchased from Janvier Labs (Le Genest-Saint-Isle, France). They were housed with a maximum of five animals per cage and had *ad libidum* access to food and water. Mice were adapted to the environment of the facility (IERP, INRAE, Jouy-en-Josas, France) one week before experiments.

Prior to infection, mice were mildly anaesthetized by intra peritoneal (i.p.) injection of ketamine (79 mg/kg) and xylazine (9 mg/kg). Bacterial strains were grown to late exponential phase and diluted in PBS. Intranasal (i.n.) bacterial administration was performed through slow instillation of 20 µL of bacterial suspension (1x 10^7^ CFU) with a micropipette (10 µL in each naris). Inoculated suspensions were plated on LB agar for CFU determination prior to injection. Empty and NM102-containing formulations were administrated in mice either: (i) *via* the i.p. route with 200 µL of suspension containing 1.5 or 6 mg/kg (1.29 and 5.16 mM, respectively) of drug or (ii) *via* the i. n. route with 20 µl of a mixture containing bacterial suspension (1x 10^7^ CFU) and NM102 formulation (1.5 or 6 mg/kg) or empty formulation. MPN (10 mg/kg) was inoculated *via* the i.p. route. For combination therapy studies, NM102 formulation and MPN were inoculated *via* the i.n. and the i.p. route, respectively. Mice were sacrified by cervical dislocation after 24 h and the left lung was placed in 2 mL sterile tubes containing 600 μL of PBS and 1.4 mm ceramic lysing beads (MP Biomedicals, Solon, OH, USA). Organs were homogenized using a FastPrep instrument (MP Biomedicals, Solon, OH, USA) and aliquots from each tube were serially diluted on LB plates for CFU enumeration. The log CFU was calculated per g of organ.

For the toxicity study, mice were treated (i.p., 200L/injection) with NM102 (6 mg/kg) or DMSO as control and mice weight was monitored for 7 days.

### Cytotoxicity

HeLa (human cervical epithelial cell line) and Vero (monkey kidney epithelial cell line) cells were cultured in Dulbecco’s Modified Eagles Medium (DMEM), supplemented with 10% heat inactivated fetal bovine serum (FBS) and 1 U/mL penicillin, and 1 µg/mL streptomycin at 37°C and 5% CO_2_. Once reaching a confluency of 70%, cells were washed with Dulbeccos Phosphate Buffered Saline (DPBS) and detached by treatment with 0,05% Trypsin-EDTA. Total cell number was determined by using a Neubauer Cell chamber and cells were seeded at a concentration of 4x10^4^ cells/well in a flat-bottom Tissue Culture 96-well plate. Cells were then incubated at 37°C and 5% CO_2_ overnight before starting the cytotoxicity assay. Two hours before the assay, medium in each well was replaced by an antibiotic-free DMEM medium. Cells were washed and treated with the respective chemical (MPN at 10 and 100 µM; NM102 at 10 and 300 µM; NM102 formulation at 1.29 and 5.16 mM) or their respective control (10% DMSO or empty formulation) in a total volume of 100 µL in Hanks’ Balanced Salt Solution (HBSS) medium (ThermoFischer). After one hour of incubation, 20 µL of CellTiter96®AQ_ueous_ was added to each well and cells were incubated again for one hour. Absorbance was measured at 490 nm after 5 seconds of shaking to determine cytotoxicity. The background of CellTiter96®AQ_ueous_ in HBSS and the background of each chemical was subtracted of the respective values. Treated cells were compared and normalized to the untreated control.

### Analysis of microbiome diversity

Mice were treated (i.p., 200 µL/injection) with DMSO or NM102 (6 mg/kg). Feces were collected (day 1 and day 7) and the quantity of bacteria was assessed by qPCR as previously described (Mayeur et al., 2013). In addition, microbiota diversity was assessed by 16S rRNA sequencing. Mice were treated (i.p., 200 µL/injection) with DMSO or NM102 (6 mg/kg) after i.n. infection with *K. pneumoniae*. Feces of untreated and treated mice were collected and stored at −80°C. Microbial DNA was extracted using the DNeasy PowerSoil Pro Kit following the provider’s instructions (Qiagen, Hilden, Germany). DNA samples were sent to the NGS Competence Center Tübingen (Tübingen, Germany) for quantification, library construction and sequencing. DNA concentration was quantified using a Qubit fluorometer (Thermo Fischer, Waltham, MA). The V4 hypervariable region of the 16S rRNA gene was amplified using primers F515 (5′-GTGCCAGCMGCCGCGGTAA-3′) and R806 (5′-GGACTACHVGGGTWTCTAAT-3′) (Caporaso et al., 2011) and sequenced with the Illumina MiSeq sequencing platform with v2 chemistry. To process 16S rRNA amplicon sequences, we used the dada2 v.1.21.0 package of R (v.4.2.0) (Callahan et al., 2016) following its standard operating procedure as available at https://benjjneb.github.io/dada2/bigdata.html. Briefly, after inspecting the quality profiles of the raw sequences, we trimmed and filtered the paired-end reads using the following parameters: trimLeft: 23, 24; truncLen: 240, 200; maxEE: 2, 2; truncQ: 11. The filtered forward and reverse reads were dereplicated separately and used for inference of the amplicon sequence variants (ASVs) with default parameters, after which they were merged in a per-sample basis. We next filtered the merged reads to retain only those with a length between 250 and 256 bp and carried out chimera removal.

We then performed the taxonomic assignment on the final set of ASVs using a curated dada2-formated database based on the genome taxonomy database (GTDB) release R06-RS202 (Parks et al., 2018). A maximum-likelihood tree of the ASVs was obtained using the R packages DECIPHER v.2.24.0 and phangorn v.2.8.1 (Schliep, 2011) after performing a multiple sequence alignment with each of the representative sequences. We used the PERFect v.1.10 package of R (Smirnova et al., 2019) to identify spurious ASVs, defined as those that made insignificant contributions to the total covariance of the microbiome dataset.

For alpha- and beta-diversity analyses, samples were rarefied to 21300 reads/sample to account for differences in library size. To compare alpha-diversity between samples, we used the Shannon index as implemented in the vegan v.2.6-2 package of R (https://CRAN.R-project.org/package=vegan). Global differences in Shannon index between treatments were assessed using the Kruskal-Wallis-Test and pairwise contrasts between treatments were performed using the Wilcoxon signed-rank test. We used the ASV-level abundance profile to obtain matrices of the weighted and unweighted UniFrac distances with the GUniFrac v.1.6 package of R (Chen et al., 2012). We assessed differences in beta-diversity estimates using the adonis function (analysis of variance using distance matrices) of the permutational multivariate analysis of variance (PERMANOVA) on the distance matrices with 9999 permutations, as implemented in the vegan v.2.6-2 package of R. For pairwise PERMANOVA tests, we used the pairwiseAdonis v.0.4 package of R (available at https://github.com/pmartinezarbizu/pairwiseAdonis). P values were adjusted for multiple comparisons using the Benjamini-Hochberg method.

### Mutation rate

*E. coli* Keio WT and Δ*mfd* strains were grown onto LB agar O/N at 37°C. Cultures in LB broth were started from a single colony at 37°C at 200 rpm until an OD _600 nm_ at around 0.005. Each culture was then divided in two with one culture exposed to 100 µM NM102 and the other to 2% final DMSO as control. Bacteria were incubated until the culture reached OD _600 nm_ of 2, which corresponds to the early stationary phase. Bacteria were plated onto LB agar plates with 50 µg/mL rifampicin and incubated for 24 h at 37°C. The number of colonies was counted and used to determine the mutation frequency. Total number of bacteria were determined following serial dilution and plating onto LB agar plate.

### Evolvability assays

A serial passage broth microdilution protocol was performed based on previously described methods (Ragheb et al., 2019) to identify the effect of NM102 on the evolution of antibiotic resistance. Briefly, culture of *E. coli* Keio WT started from a single colony was diluted to OD _600 nm_ = 0.005. Bacterial suspension was subsequently transferred into a 96-well microplate (180 μL/well) and incubated with NM102, DMSO 2% or left untreated. Treated and untreated bacteria were grown in LB with a gradient of concentrations of the indicated antibiotic to select for resistance for 18 h-24 h at 37°C under stationary conditions. The OD _600 nm_ was measured and MICs were determined. Serial passage was performed using cells growing (defined by an OD _600 nm_ at a minimum of 50% of the untreated condition) in the highest concentration of antibiotic. Antimicrobial concentrations were adjusted during the process to compensate for the rise in MIC values. This process was repeated for 4 or 5 passages. Results were expressed as the median MIC of the 3 replicates.

---

**Supp Fig. 1.**
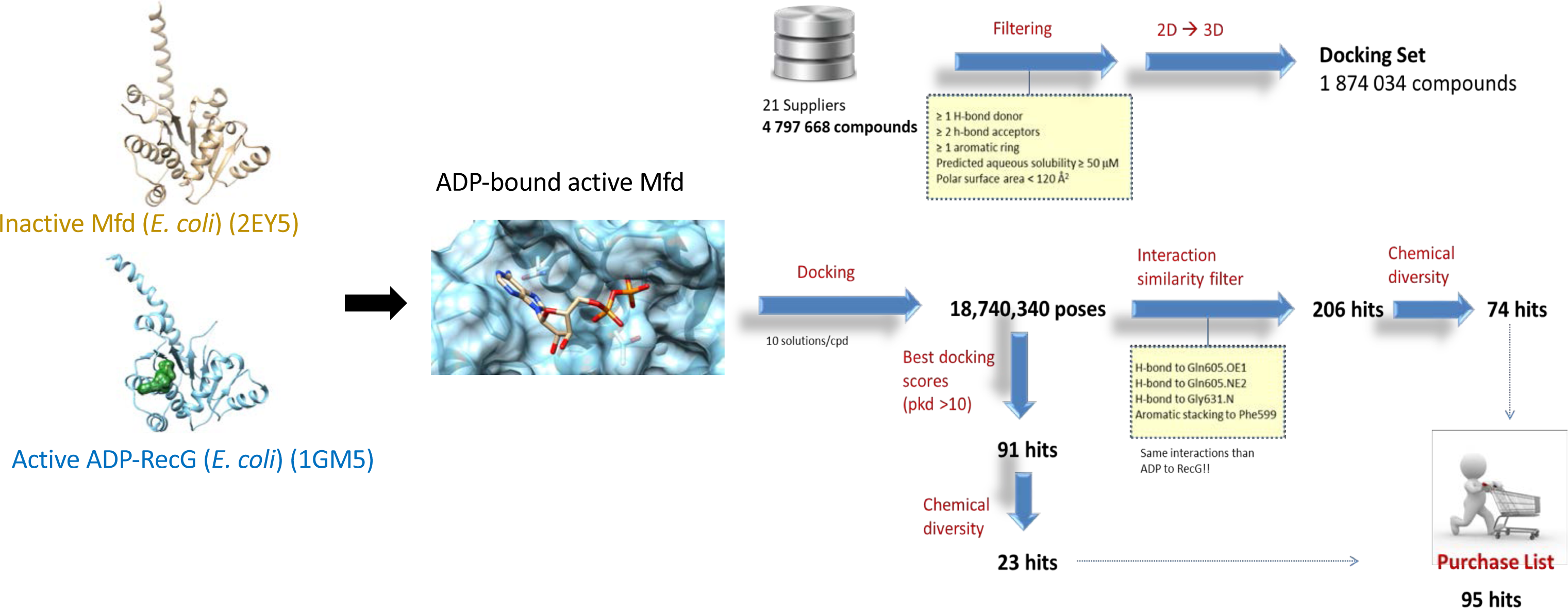
Modelling of active Mfd of *E. coli* and high throughput screening of compounds library. Left: 3D model of active *E. coli* Mfd allows the docking of ADP and molecules. Right: Virtual screening workflow to select 95 commercially-available potential Mfd inhibitors.

**Supp Fig 2.**
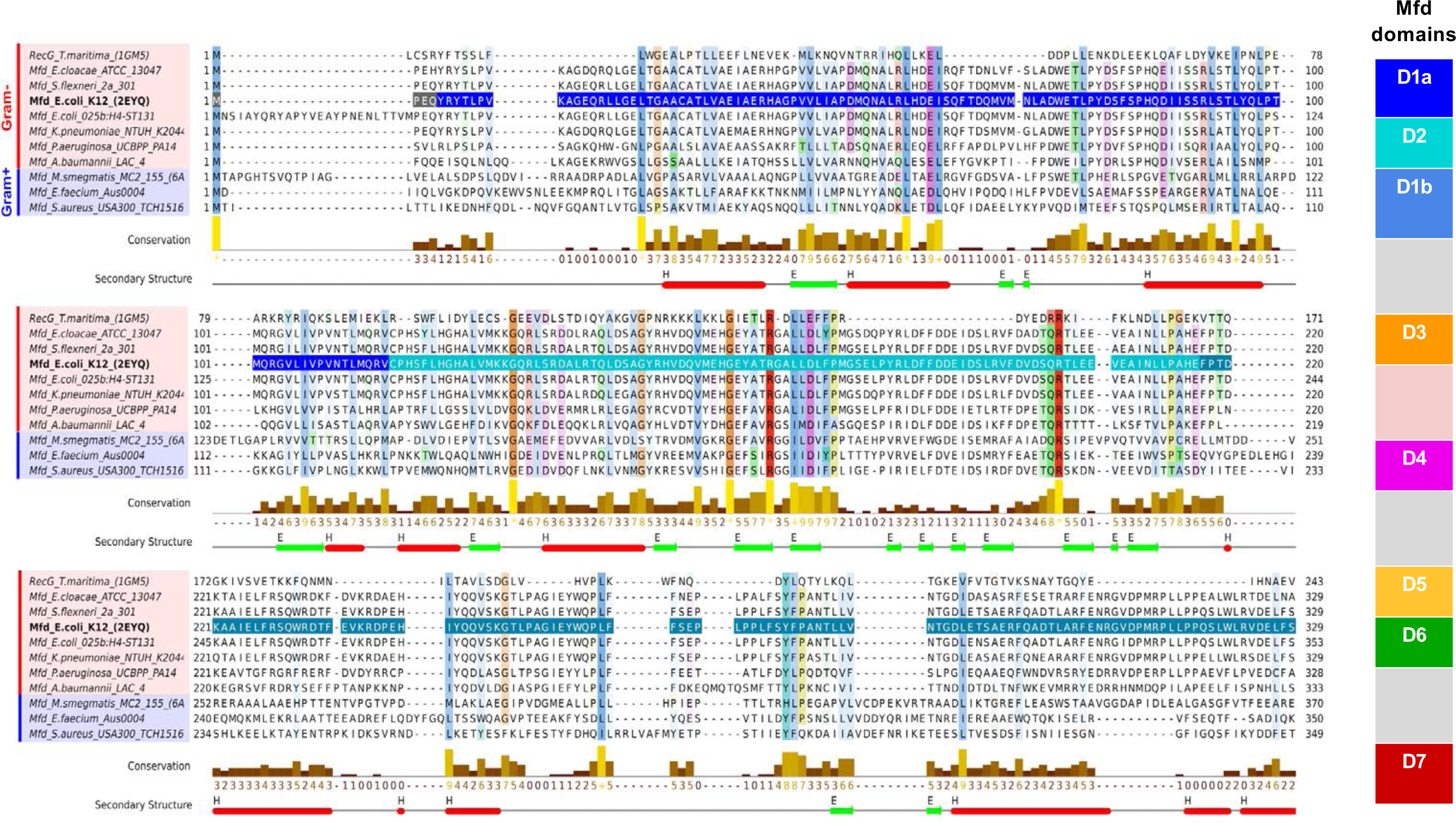
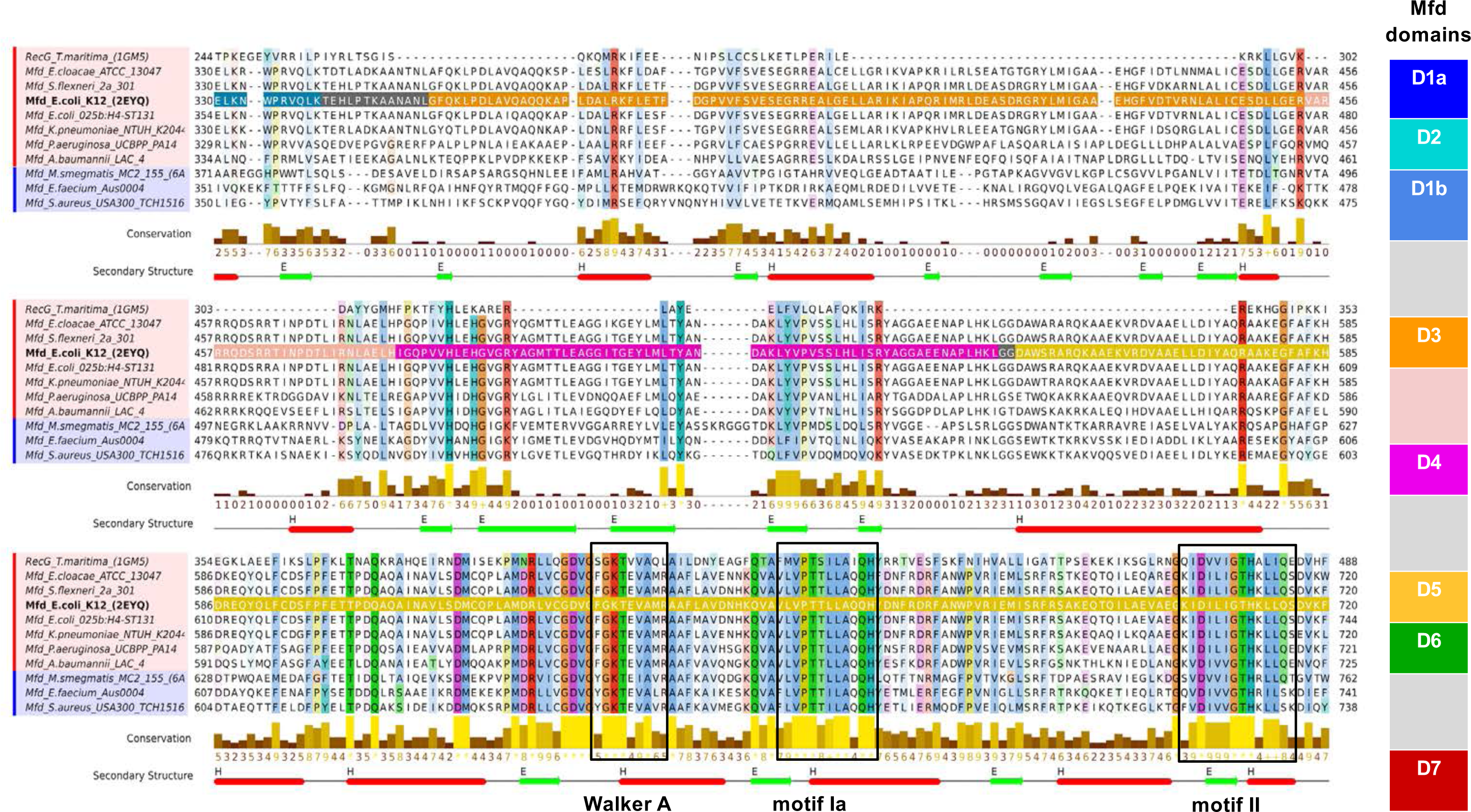
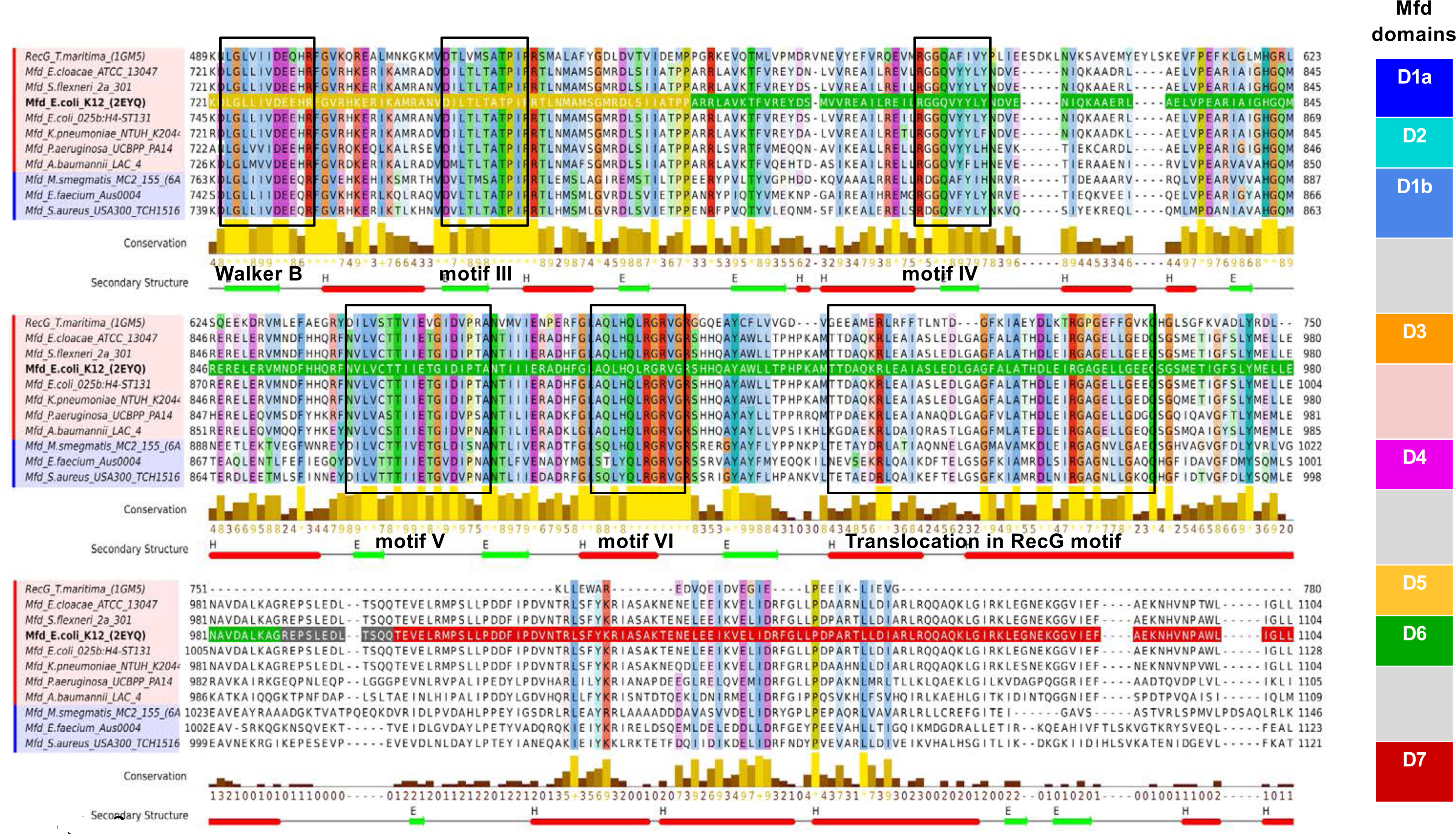
Mfd alignment. Multiple sequence alignment of Mfd from ESKAPE and RecG from *T. maritima* using MAFFT. The vertical stripe gives the color code of the multiple domains of Mfd. This color code is highlighted in the multiple alignment sequence along Mfd*_E. coli_* sequence. The typical functional motifs of an ATP-ase protein exclusively located in domains D5 and D6 are annotated and highlighted in black.

**Supp Fig 3.**
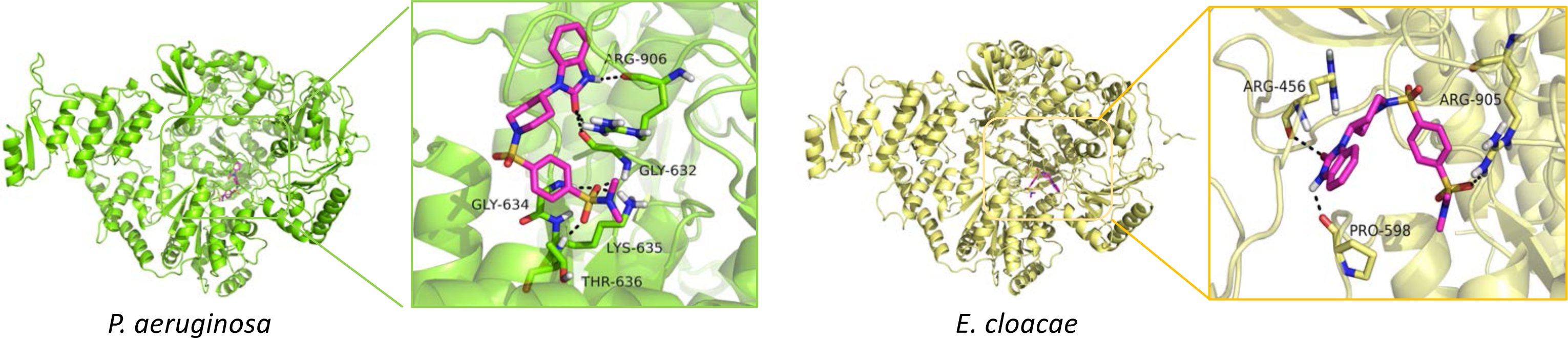
Homology models of Mfd from ESKAPE bacteria and NM102 docking. Inserted are close-views of their respective active site with residues involved in the binding. The NM102 is shown as pink stick.

**Supp Fig.4.**
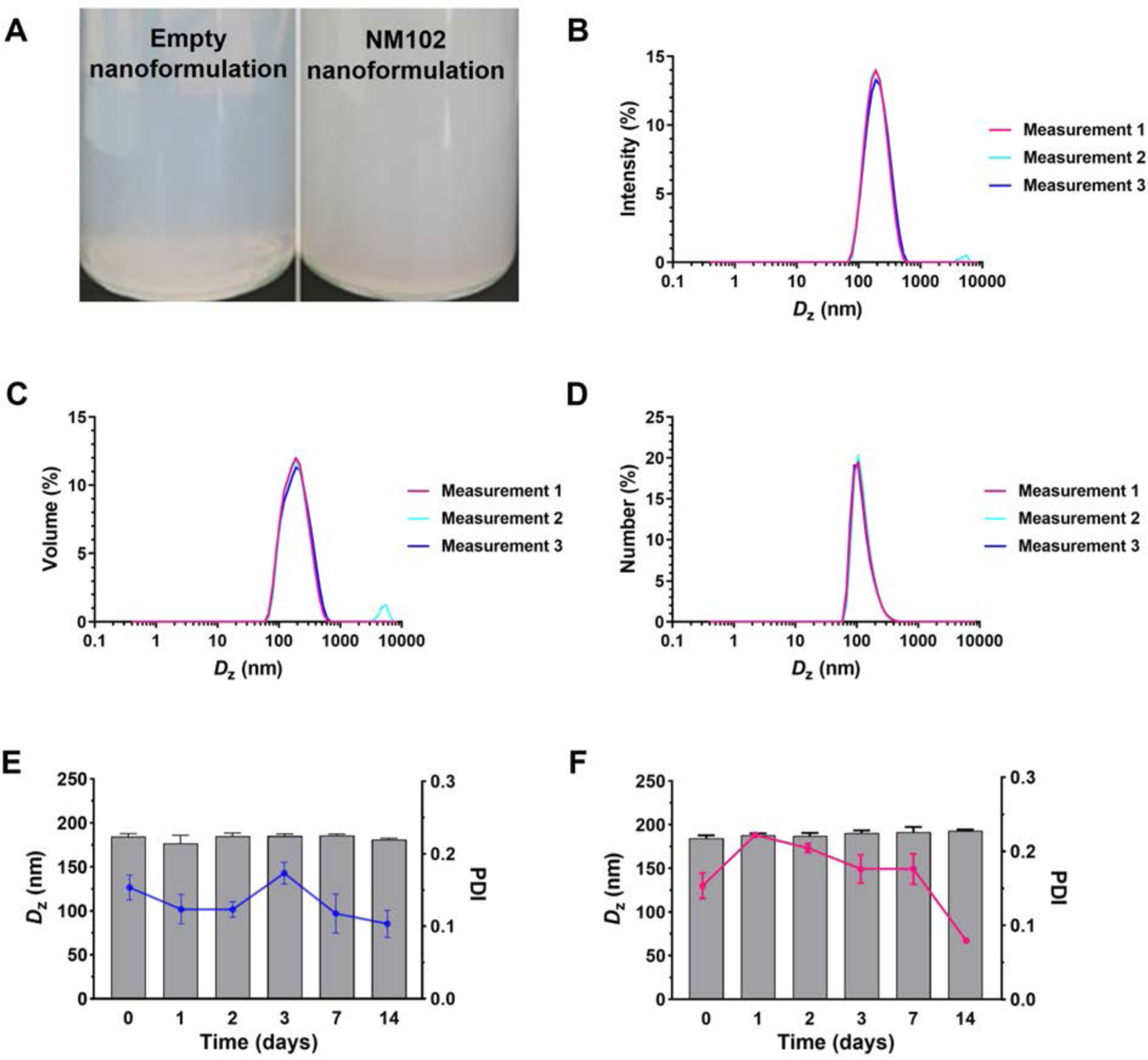
NM102 nanoformulation. (A) Represenative images of empty and NM102 nanoformulations. Size distribution of NM102 nanoformulation by (B) intensity, (C) volume and (D) number. Evolution with time of intensity-averaged diameter of NM102 nanoformulation at (E) 4°C and (F) room temperature up to 14 days.

**Supp Fig.5.**
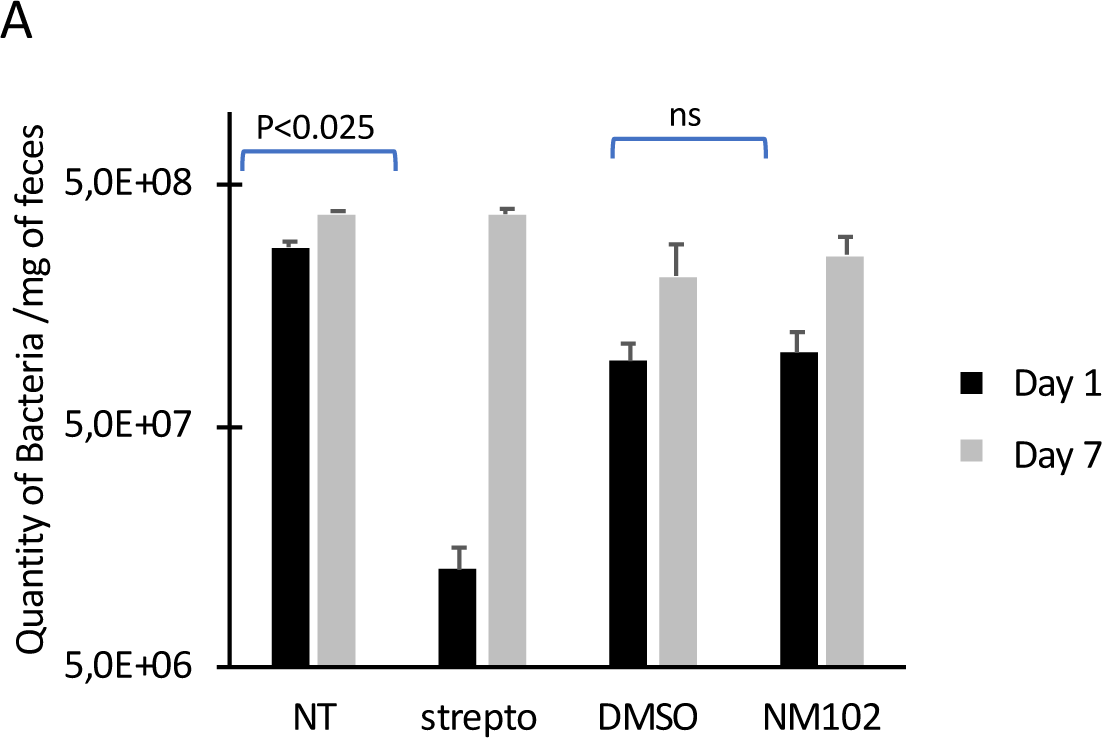
NM102 does not impact the host microbiome. The quantity of total bacteria present in the mice feces was quantified by qPCR at day 1 and day 7 post-administration of NM102. The graph shows the mean ± SD of two independent samples.

**Supp Fig.6.**
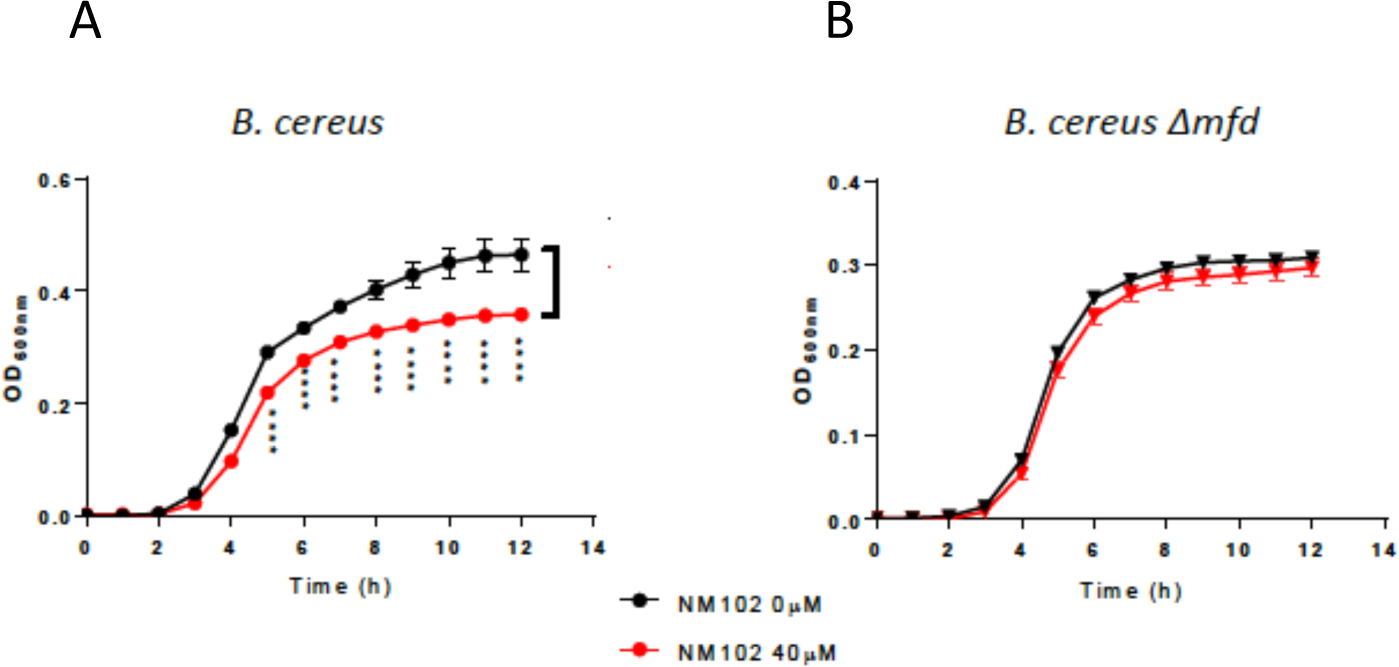
NM102 decreases *B. cereus* survival in the context of NO-stress. *B. cereus* wt and Δ*mfd* strains were exposed to NM102 (40 µM) and NOC 5 and bacterial growth was followed by measuring the OD_600 nm_ for 12 h. The results reported are mean ± SD of three independent experiments each in triplicates, P values are calculated against the condition without NM102, using One Way ANOVA (**** P<0.0001).

**Supp Table 1.** Table of conserved 3D positions involved in the binding of NM102 for the ESKAPE group. Residues hydrophobic, acidic, polar, basic and glycine are colored in green, salmon, cyan, wheat and marine, respectively. Stars highlight residues which are not strictly conserved.

|  | P1 | P2 | P3 | P4 | P5 | P6 | P7 | P8 | P9 | P10 | P11 | P12 | P13 | P14 | P15 | P16 | P17 | P18 | P19 | P20 | P21 |
| --- | --- | --- | --- | --- | --- | --- | --- | --- | --- | --- | --- | --- | --- | --- | --- | --- | --- | --- | --- | --- | --- |
| *E.coli* | F597 | F599 | E600 | T602 | Q605 | D629 | V630 | G631 | F632 | G633 | K634 | T635 | E636 | Q664 | H665 | N668 | E730 | P780 | A781 | D876 | R905 |
| *E. faecium* | F618 | Y620 | S621 | T623 | Q626 | D650 | V651 | G652 | Y653 | G654 | K655 | T656 | E657 | Q685 | H686 | T689 | E751 | P801 | A802 | D897 | R926 |
| *S.aureus* | F615 | Y617 | E618 | T620 | Q623 | D647 | V648 | G649 | Y650 | G651 | K652 | T653 | E654 | Q682 | H683 | T686 | E748 | P798 | E799 | D894 | R923 |
| *K.pneumoniae* | F597 | F599 | E600 | T602 | Q605 | D629 | V630 | G631 | F632 | G633 | K634 | T635 | E636 | Q664 | H665 | N668 | E730 | P780 | A781 | D876 | R905 |
| *A. baumannii* | F602 | Y604 | E605 | E606 | T607 | D634 | V635 | G636 | F637 | G638 | K639 | T640 | E641 | Q669 | H670 | S673 | E735 | P785 | A786 | D881 | R910 |
| *P.aeruginosa* | F598 | F600 | E601 | E602 | T603 | D630 | V631 | G632 | F633 | G634 | K635 | T636 | E637 | Q665 | H666 | N668 | E731 | P781 | A782 | D877 | R906 |
| *E. cloacae* | F597 | F599 | E600 | T602 | Q605 | D629 | V630 | G631 | F632 | G633 | K634 | T635 | E636 | Q664 | H665 | N668 | E730 | P780 | A781 | D876 | R905 |
| *S.flexneri* | F597 | F599 | E600 | T602 | Q605 | D629 | V630 | G631 | F632 | G633 | K634 | T635 | E636 | Q664 | H665 | N668 | E730 | P780 | A781 | D876 | R905 |
|  |  | * | * | * |  |  |  |  | * |  |  |  |  |  |  | * |  |  | * |  |  |

**Supp Table 2:** ESKAPE strains used in this study and their resistance to antibiotics. The MIC for each antibiotic tested is in µg/mL.

|  | **K. pneumoniae**<br>**ATCC 700603** | **K. pneumoniae**<br>**ALE CTXM-15+** | **K. pneumoniae**<br>**DOU CTXM-15** | **P. aeruginosa**<br>**CIP 27853** | **E. coli**<br>**ATCC 25922** |
| --- | --- | --- | --- | --- | --- |
| **Penicillin** | 16 | >512 | >512 | 32 | 1 |
| **Ampicillin** | 8 | 4 | >512 | 64 | 4 |
| **Ceftriaxone** | >512 | >512 | >512 | >512 | 16 |
| **Meropenem** | 2 | >512 | >512 | 16 | <0.5 |
| **Cefotaxin** | <0.5 | 8 | 8 | <0.5 | <0.5 |
| **Oxacillin** | 8 | >512 | >512 | 16 | <0.5 |
| **Streptomycin** | >512 | >512 | >512 | >512 | 2 |
| **Ciprofloxacin** | >512 | >512 | >512 | >512 | 4 |

**Supp Table 3.** The solubility of NM102 in some common solvents.

| Excipient | Maximal solubility of NM102 after 24 h at 20°C, quantified by HPLC (mg/mL) |
| --- | --- |
| Water | $\leq 0.001$ |
| Tween 80 | 0.09 |
| Intralipid 20% | 0.05 |
| Soya oil | $< 0.002$ |
| Corn oil | $< 0.002$ |
| Capmul® MCM | 0.01 |
| Mygliol 912 | 0.002 |
| NMP | $\geq 10$ |
| DMSO | $\approx 6.7$ |

